# G4 DNA structures induced by UV radiation: a multi-omic approach

**DOI:** 10.64898/2026.07.15.738736

**Authors:** Lindsay R. Julio, Diana V. Turrieta Vejar, Charlotte H. Walborsky, Annika Salpukas, Ella Chee, Julianne C. Murthy, Shannon G. MacLeod, Adrianna L. Vandeuren, Mynaja Ferguson, Rachel E. Muriph, Jason J. Evans, Tovah A. Day

## Abstract

DNA G quadruplexes (G4s) are secondary structures with critical roles in regulating genome function. G4s in regulatory regions like promoters and enhancers are important for controlling gene expression, while aberrant formation of G4s has been linked to genomic instability and human disease. Despite G4s’ importance and inherent danger, their dynamic formation remains incompletely understood. Here, we show that ultraviolet (UV) radiation uniquely induces widespread and persistent G4 formation in human cells, distinguishing it from other genotoxins. Through integrated genomic, transcriptomic, and proteomic analyses, we uncovered key features and functions of UV-induced G4s. Proteomic profiling identified RCOR3 as a factor associated with specific UV-induced G4s at late time points. Functional studies revealed that RCOR3 is essential for the formation and persistence of these structures. Furthermore, genes associated with UV G4s that are differentially expressed are enriched in pathways related to response to UV radiation, highlighting their biological relevance. These findings define the multi-omic landscape of UV-induced G4s and reveal new mechanistic insights into the interplay between genotoxic stress responses and the regulation of non-canonical DNA structures.

**GRAPHICAL ABSTRACT:** 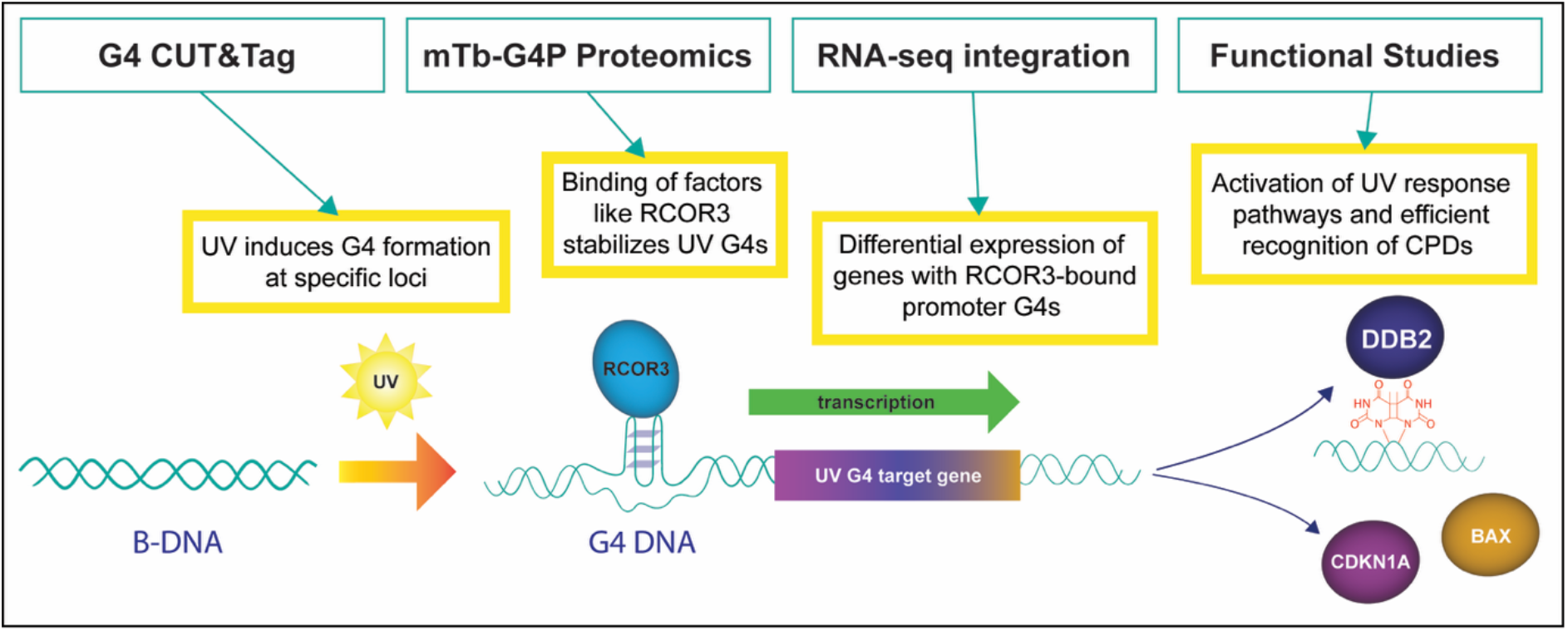

## INTRODUCTION

Ultraviolet (UV) radiation is a ubiquitous environmental mutagen that poses a significant threat to human health in the form of photodamage, photoaging, and skin cancer. When UV-B radiation, the primary driver of sunburn and skin cancer, reaches DNA molecules, it causes dimeric photoproducts primarily in the form of cyclobutane pyrimidine dimers (CPDs) and pyrimidine-pyrimidone (6–4) photoproducts. These DNA lesions are repaired by nucleotide excision repair (NER) pathways; a critical step in NER is the recognition of the DNA lesion by the DDB1/DDB2 dimer; upon recognition, the lesion is removed and DNA is resynthesized (1). Failure to repair UV-induced DNA lesions leads to the accumulation of mutations which drive the development of skin cancer and patients with germline mutations in NER factors have high rates of skin cancers resulting from their inability to repair these lesions. (2). UV exposure can also trigger inflammation (3), vascular remodeling (4), checkpoint activation (5), and even apoptosis to prevent cells with damaged genomes from replicating (6,7).

Decades of research have characterized UV-induced DNA lesions, yet emerging evidence points to a distinct and persistent structural consequence of UV exposure: the dynamic conversion of B DNA into G quadruplex (G4) DNA, a non-canonical four-stranded DNA structure. This B-to-G4 conversion has been observed at early timepoints following UV in both yeast and human cells (8,9). These conversions are possible in guanine-rich sequences, where non-consecutive guanines can assemble by Hoogsteen base pairing into planar G-quartets that stack into a stable, four-stranded G4 conformation. Computational and deep-sequencing analyses have identified more than 700,000 sequences in the human genome with the potential to form intramolecular G4s (10–12). However, only about 1% fold into G4 structures within cells (13,14), and the specific G4s that form differ substantially across cell types (15). Together, these findings indicate that G4 formation is tightly regulated rather than merely determined by sequence alone. G4 DNA structures are known to regulate transcription (16,17), DNA replication (18,19), and telomere maintenance (20,21). In particular, in human cells the majority of G4 structures are located at gene regulatory elements, including transcription start sites, promoters, and enhancers (17), and promoter G4s are typically associated with increased transcriptional activity (15–17,22). G4s also promote transcription by recruiting transcription factors (17,23) and chromatin-modifying enzymes (14,16,24) to their genomic sites.

Recently, several observations have suggested that G4 structures interact with DNA repair pathways by multiple mechanisms (25). For example, two central members of the NER pathway, the helicases XPD and XPB (26), preferentially bind sequences containing G4 motifs (27) with their archaeal homologs interacting directly with G4 structures *in vitro* (27). In yeast, NER components Rad23, Rad1, and Rad2, the homologs of human XPC, XPF, and XPG, bind G4s stabilized by Zuo1 (human ZRF1) (8). Notably, yeast lacking Zuo1 are UV-sensitive, and this sensitivity can be rescued by small molecule stabilization of G4s (8), suggesting that G4s are contributing positively to the cellular response to UV. In addition, G4s have been suggested to promote template strand invasion during homologous recombination, thereby facilitating the repair of DNA double-strand breaks (28). Finally, the guanines in G4 motifs are particularly susceptible to DNA damage by reactive oxygen species (ROS) (29). Repair of the oxidized guanine (OG) occurs by base excision repair (BER); the OG is first removed to create an apurinic site which recruits the enzyme APE1. This sequence of steps leads to G4 folding and a subsequent increase in gene regulation (30–32), suggesting that genotoxin exposure, in this case ROS, can lead to G4-dependent increases in gene expression. The details of additional mechanisms of crosstalk between G4 DNA and DNA damage remain to be investigated.

Here, we propose an additional dimension to the G4-DNA damage crosstalk: UV radiation induces persistent G4 structures at specific loci throughout the human genome. These structure, in turn, drive a transcriptional program to support efficient repair of UV lesions. We identify a group of proteins that are important for UV-induced G4 formation and focus upon RCOR3, an under characterized homolog of the REST corepressor 1 (RCOR1) protein. Upon UV exposure, RCOR3 binds and stabilizes UV G4 loci identified by <u>C</u>leavage <u>U</u>nder <u>T</u>argets and <u>Tag</u>mentation (CUT&Tag), including at the promoter of NER recognition factor DDB2. In the absence of RCOR3, UV G4 induction is abrogated, increases in gene expression associated with UV G4s are lost, and cells fail to process UV-induced DNA lesions in an efficient manner.

## MATERIAL AND METHODS

### Cell Culture

A549, A375, and 293T cells were cultured in DMEM (Gibco) supplemented with 10% fetal bovine serum (FBS), and 50 U ml^−1^ penicillin and 50 ng/mL streptomycin (Gibco). Cells were tested for mycoplasma contamination using the Lonza Mycoplasma kit. All cells were cultured at 37^ο^C under 5% CO_2_ atmosphere.

#### UV Irradiation of cells

Cells were plated at ∼25% confluence 3 days before harvest or fixation. At given timepoint pre-harvest or fixation, media was aspirated from cells and set aside. Warm PBS was added to cover cells. Cells were irradiated with 10J/cm^2^ of 302 nm UVB in a CL-3000 UV irradiator (Analytik Jena). PBS was removed and media was replaced.

#### siRNA Transfection

ZRF1 and RCOR3 ON-TARGETplus Human Smartpool siRNAs were obtained from Horizon Discovery (previously Dharmacon, L-017112-02 and L-025435-01). siRNA pools for ASCC2, XRCC4, SAP30BP, UFD1 (UFD1L), CCDC9, USF1, KAT6B (MORF), ARPC1B, KAT7 (HBO1), CUL1, KMT2C (MLL3), and SUV39H2 (KMT1B) were purchased from IDT. Cells were transfected with 86 nM siRNA using Lipofectamine 3000 transfection reagent (Fisher Scientific, L3000015) for 48 hours. The siRNA sequences are listed in (**Table S2**).

#### Immunofluorescence microscopy (BG4 and HA tag)

Cells were plated on Nunc™ Lab-Tek™ II CC2™ 8-well slides (ThermoFisher 154941) three days prior to fixation and irradiated as indicated. Where indicated, siRNA transfection or doxycycline (2 μg/mL) induction of Tet-on promoters was carried out at 48 hr prior to fixation. At the designated times after UV treatment, doxycycline induction, or transfection, cells were washed twice with ice cold PBS and fixed for 10 minutes in 4% PFA in PBS. Cells were permeabilized in PBS-Triton for 10 minutes at room temperature and RNA was degraded with 0.24 mg/mL RNase A in PBS (NEB) for 1 hour at 37^ο^C. Cells were blocked with 0.5% goat serum in PBST (PBS + 0.1% Tween-20) for 1 hr at 37°C and incubated with rabbit anti-HA (Bethyl A190-108A) at 1:1500 dilution in blocking buffer for 1 hour or flag-tagged BG4 sc-Fv (in-house) at 1:100 dilution for 1 hour at 37^ο^C followed by a rabbit-anti-flag secondary antibody (Cell Signaling Technology, 2368S) at 1:800 dilution for 1 hour at 37^ο^C. In both cases, Alexa Fluor 594 Goat anti-Rabbit antibody (ThermoFisher, A-11012) was used at 1:1000 dilution for 1 hour at room temperature. Cells were washed thrice in PBST after RNase treatment and between each antibody incubation. Cells were mounted using VectaShield Anti-Fade mounting medium with DAPI (Fisher Scientific, NC1695563) and imaged at 40x magnification on a Zeiss 880 confocal microscope.

### Plasmids

#### MiniTurbo plasmids

MiniTurbo fragment was obtained by PCR from the plasmid 3xHA-miniTurbo-NLS_pCDNA3 (Addgene #107172, a gift from Alice Ting; https://www.addgene.org/107172/) (33). Codon optimized G4P fragment was generated by ThermoFisher and a PCR product containing attB sites was used to generate Multi-Site Gateway (ThermoFisher) entry clones by BP reaction (**Table S3**). Entry clones were subsequently recombined into inducible lentiviral expression vector pCW57.1 (Addgene #41393, a gift from David Root; https://www.addgene.org/41393/) by Multi-Site Gateway LR reaction.

#### CRISPRi plasmids

gRNA oligonucleotides (**Table S4**) were cloned into lentiGuide-Puro lentiviral expression vector (Addgene #52963, a gift from Feng Zhang; https://www.addgene.org/52963/) by Golden Gate assembly.

### Lentivirus

#### Preparation of lentivirus

Guide plasmid constructs, CRISPRi construct pHR-UCOE-SFFV-Zim3-dCas9-P2A-Hygro, (Addgene #154472, a gift from Mikko Taipale; https://www.addgene.org/154472/) (34), or pCW57.1 constructs containing mTb-G4P or mTb alone, were cotransfected into LentiX 293T cells (Takara) alongside packaging plasmid (psPAX2, Addgene #12260; https://www.addgene.org/12260/) and envelope plasmid (pMD2.G, Addgene #12259; https://www.addgene.org/12259/) using Lipofectamine 3000 (Invitrogen). After transfection, cells were cultured in in DMEM (Gibco) supplemented with 30% fetal bovine serum (FBS), and 50 U /mL penicillin and 50 ng/mL streptomycin (Gibco). Both psPAX2 and pMD2.G were gifts from Didier Trono.

#### Lentiviral transductions

100 μL of lentivirus and polybrene (final concentration of 10 μg/mL) were added to subconfluent LentiX cells in 6-well plates. Cells were spun at 1,178 x *g* for 15 min at RT and returned to incubator overnight. Twenty-four hours after transduction, cells were cultured in selection medium. Puromycin (Gibco) selection for miniTurbo constructs in A549 and A375 cells was at 1.0 μg/mL for 48 hours. Hygromycin B (Invitrogen) selection for CRISPRi construct in A375 cells was at concentration 200 μg/mL for 8 days, and Blasticidin (Sigma) selection for guide RNA constructs in A375 cells was at concentration (10 μg/mL) for 5 days.

### BG4 production

The BG4 production was adapted from the protocol described by De Magis *et al.* (35). Briefly, the pSANG10-3F-BG4 plasmid, (Addgene, #55756, a gift from Shankar Balasubramanian; https://www.addgene.org/55756/ (36)), was transformed in BL21(DE3) competent cells (Thermo Scientific, EC0114). Transformed cells were grown in 2L YT media (Sigma-Aldrich, Y2377-250G) with 50 μg/mL Kanamycin (Fisher Scientific, BP906-5) at 37°C at 220 RPM for approximately 9 hours. BG4 antibody expression was induced by adding 0.5 mM Isopropyl-β-D-thiogalactopyranoside (MP Biomedicals, 114064112) and incubating overnight at 25°C. Bacteria were harvested by centrifugation at 4,500x*g* at 4°C for 12 min and lysed in TES buffer (50mM Tris-Cl pH 8.0, 1mM EDTA, 20% sucrose) on ice for 10 min. The lysate was spun at 16,000x*g* at 4°C for 1hr and the supernatant was filtered through a 0.45 μm vacuum filter. BG4 protein was then purified on a Ni-NTA Sepharose column (ThermoFisher, 88226) pre-equilibrated with TES buffer. The column was washed twice with PBS supplemented with 100nM NaCl and 10mM Imidazole (pH 8.0) by centrifugation at 700x*g* at 4°C for 2 min. BG4 was eluted in PBS with 250 mM Imidazole and EDTA-free cOmplete Protease Inhibitor Cocktail (Roche, 11873580001). The elution buffer was then exchanged by inner cell salt buffer (25mM HEPES pH 7.6, 110mM KCl, 10.5mM NaCl, 1mM MgCl_2_ by overnight dialysis (ThermoFisher, 66830) and subsequently the BG4 was concentrated using the Amicon Ultra-15 centrifugal Filter unit with 10 kDa cutoff (Millipore Sigma, UFC901024). The purity and concentration of the BG4 preparation was determined by SDS-PAGE.

### G4 measurement in mouse tissues

#### UV irradiation of mouse tissue

Animals were sedated with intraperitoneal ketamine/xylazine anesthesia (50 mg/kg ketamine; 5 mg/kg xylazine) and shaved with clippers prior to UVB exposure. Animals were exposed to a UVB dose of 20 J/cm^2^ using a dosimeter calibrated UV instrument (310 nm peak output) (Tyler Research, Alberta, Canada). Full-thickness dorsal skin samples in telogen phase were collected and fixed at 24- and 48 hr after irradiation in addition to sham controls and paraffin embedded cross-sections were prepared for histologic analysis. Animals were euthanized by cervical dislocation.

#### Immunohistochemistry

Tissues were fixed overnight in 10% neutral buffered formalin and thereafter transferred to 70% ethanol at 4°C until processing. Tissues were processed using an automated tissue processor (Thermo HistoStar, Kalamazoo, Michigan, USA) and embedded in paraffin wax. A microtome (Leica Biosystems, Buffalo Grove, Illinois, USA) was used to cut 4 μm skin cross-sections, which were allowed to dry overnight before de-waxing and further processing. Heat induced epitope retrieval was performed on de-waxed slides with either citrate or Tris-EDTA buffer prior to immunofluorescence staining. Skin cross-sections were blocked for 30 minutes at room temperature in 5% normal goat serum PBS, which was also the primary antibody diluent. Primary antibodies and their usage were as follows: 1:100 BG4 (EDM Millipore, MABE917) in blocking buffer at room temperature for 2 hours. This was followed by incubation with anti-FLAG secondary (1:200, Cell Signaling Technology, 2368S) and Alexa Fluor 488 tertiary (1:200, ThermoFisher, A-11012) for 30 minutes each at room temperature. Slides were mounted with ProLong Gold mounting media with DAPI (4’,6-diamidino-2-phenylindole, (Invitrogen P36931) and allowed to cure prior to imaging on a Zeiss 710 confocal microscope.

### Image analysis

FIJI ImageJ (37) was used to quantify BG4 foci and intensity per nucleus or HA intensity per nucleus for all immunofluorescence and immunohistochemistry experiments. Images from the vehicle-treated condition were used to set an appropriate noise tolerance for each antibody stain.

### Biotin labelling with miniTurbo

#### Biotinylation

A375 cells were seeded at ∼25% confluence in 2x 15 cm^2^ dishes per condition 3 days prior to harvest. UV irradiation was carried out at timepoints as indicated. Fusion protein construct expression was induced by 2 μg/mL of doxycycline (ThermoFisher) at 48 hr prior to fixation. 30 minutes prior to harvest, media was replaced with fresh DMEM supplemented with 10% FBS, 1% penicillin/streptomycin and 500 μM exogenous biotin and incubated at 37^ο^C under 5% CO_2_ atmosphere.

#### Nuclear extraction of protein

Nuclear extraction and immunoprecipitation of biotinylated protein was adapted from the protocols described by Yan *et al.* (38) and Cho *et al.* (39). Cells were washed 2x with warm PBS and trypsinized. Pellets of 20 million cells per condition were further washed 3x with ice cold PBS. Cells were resuspended in Extraction Buffer A (10 mM HEPES pH 7.9, 5 mM MgCl_2_, 0.25 M sucrose, 0.1% NP-40) and incubated on ice for 10 minutes. Nuclei were pelleted by centrifugation at 6,000x*g* and 4^ο^C for 10 minutes, resuspended in Extraction Buffer B (10 mM Hepes pH 7.9, 1.5 mM MgCl_2_, 0.1 mM EDTA and 25% glycerol, 0.42 M NaCl), and incubated on ice for 20 minutes. Lysate was spun down for 15 minutes at 9,400x*g* at 4^ο^C for 15 minutes, at which point supernatant was transferred to new tubes and nuclear protein concentration was measured by Qubit BR Protein Assay (ThermoFisher). Protein aliquots were set aside for western.

#### Immunoprecipitation of biotinylated proteins

Streptavidin magnetic beads (ThermoFisher, 88817) were washed with TBS-Tween (25 mM Tris-HCl, pH 7.2, 0.15 M NaCl, 0.1% Tween-20) and 50 μL of washed beads were added to 150 μg of nuclear protein and incubated overnight at 4^ο^C with end-over-end rotation. Beads were washed 2x for 5 minutes each with 1% SDS, 2x for 5 minutes with BC500 buffer (50 mM Tris-HCl pH 7.6, 2 mM EDTA, 500 mM KCl), 1x with BC100 buffer (50 mM Tris-HCl pH 7.6, 2 mM EDTA, 100 mM KCl) with 2 M Urea and 2x for 5 minutes in BC100 (without Urea).

### Mass Spectrometry

Pooled technical replicates were subjected to 1D LC-IM-MS using a timsTOF HT mass spectrometer (Bruker) utilizing trapped ion mobility separation and data-independent acquisition. Samples were run against a 1% false discovery rate. Mass spectrometry was performed at the Proteomics Core Facility at the University of Massachusetts Boston.

### Western blotting

#### Protein extraction

Cells were harvested and lysed with RIPA buffer (ThermoFisher, 89901) containing protease inhibitor (ThermoFisher, 78430) for 5 min on ice. They were subsequently sonicated for 4 cycles (30 sec on, 30 sec off) using the Bioruptor Pico (Diagenode) and centrifuged for 15 min at 14,000x*g*.

#### Blotting

30μg protein per sample was denatured in Laemmli buffer at 95^ο^C for 5 minutes. Samples were resolved on a 10% polyacrylamide gel run 30min at 50V and 1h at 150V. Proteins were then transferred to a nitrocellulose membrane (Bio-Rad, 1620112) using a semi-dry transfer apparatus (Bio-Rad) run at 16V for 1.5hr and 10V for 45 min. Blots were blocked using 5% milk in PBST (0.05% Tween-20) for 45 min, then probed with primary antibody in blocking buffer. Primary antibodies were as follows: anti-ZRF1 Ab (1:1000; clone D6B1E, CST #12844), anti-RCOR3 Ab (1:1000; Abcam ab76921), anti-P53 Ab (1:1000; CST #9282), anti-HA Ab (Bethyl A190-108A). Blots were washed 3 times with PBST and incubated with goat-anti-rabbit IgG StarBright Blue700 secondary antibody (1:2500; Bio-Rad 12004161) for 1 hr. Biotinylation of protein extracts was visualized by incubating with Streptavidin-IRDye680 (1:2500, LICOR 926-68079). for 1 hr at RT. Blots were imaged on the ChemiDoc Gel Imager (Bio-Rad), band intensities were quantified using FIJI ImageJ software (37), and the protein levels were normalized using anti-Actin hFAB Rhodamine Ab (1:2500; Bio-Rad, #12004163).

### Dot Blot

Genomic DNA was extracted using the Monarch Spin gDNA extraction kit (NEB). 150 ng DNA per condition was diluted in water and loaded onto a positively charged nylon membrane (Sigma) using the Bio-Dot Microfiltration apparatus (BioRad). DNA was then crosslinked in an oven at 80 °C for 1.5 h, blocked in 2% bovine serum albumin in PBST for 1 hr at RT, and incubated in primary antibody overnight at 4°C (anti-thymidine dimer mAb 1:500, clone KTM53, Kamiya Biosciences MC-062). Blot was incubated in secondary detection antibody conjugated with AlexaFluor-488 (Thermo Fisher, A-11012) for 1 hr and imaged on the ChemiDoc Gel Imager (Bio-Rad). Dot intensities were normalized to SYBR Gold (Invitrogen) after 48 hr incubation in 1x dilution for 48 hr.

### G4 CUT&Tag

The CUT&Tag protocol was adapted from Lyu *et al.*, 2022 (40) with modifications. Briefly, 5×10^5^ cells were harvested per condition, washed with PBS and resuspended in NP40-digitonin buffer (20 mM HEPES, pH 7.5, 150 mM NaCl, 0.5 mM Spermidine, 0.05% digitonin, 0.01% NP40 and Roche cOmplete Protease Inhibitor EDTA-free). The cells were then centrifuged at 600x*g* for 3 min, resuspended in 200 μL antibody buffer (NP40-digitonin buffer with 1% BSA and 2 mM EDTA) containing 2 μg of BG4 (purified from *E. coli*), and 35 ng Drosophila Spike-in Chromatin (Active Motif, 53083) and incubated overnight at 4°C. Nuclei were washed once with NP40-digitonin buffer followed by centrifugation at 600x*g* for 4 min, then resuspended in 200 μL antibody buffer containing 1:50 anti-Flag Ab (Sigma-Aldrich, F1804-200UG) and incubated for 1 hr at room temperature with end-over-end rotation. Samples were washed 3x and resuspended in 200 μL antibody buffer with 1:50 anti-mouse Ab (Sigma-Aldrich, M7023) and incubating for 1 hr at room temperature. The nuclei were then washed 3x and resuspended in 200 μL dig-300 wash buffer (20 mM HEPES, pH 7.5, 300 mM NaCl, 0.5 mM Spermidine, 0.01% digitonin, 0.01% NP40 and Roche cOmplete Protease Inhibitor EDTA-free) with 2.5 μL pA-Tn5 (EpiCypher, 15-1017) and incubated at room temperature. After three washes with dig-300 wash buffer with centrifugation at 300x*g* for 6 min, the nuclei were resuspended in 200 μL tagmentation buffer (dig-300 wash buffer containing 10 mM MgCl_2_) and incubated at 37°C for 1hr. The reaction was stopped by adding 6.7 μL of 0.5 M EDTA, 2 μL of 10% SDS and 5 μL of 20 mg/mL proteinase K and incubating at 63°C for 1 hr. DNA was purified using a DNA Clean & Concentrator-5 kit (Zymo, D4014), eluted in 20 μL, and RNA was digested by adding 1 μL RNase cocktail (Fisher Scientific, AM2288) with incubation for 20 min at 37°C. For library prep, 2 μL of 10 μM uniquely barcoded universal i5 and i7 primers (**Table S5**) and 25 μL NEBNext Ultra II Q5 2x PCR Master mix (NEB, M0544L) were added to each 21 μL sample and a short PCR was performed (72°C for 5 min, 98°C for 30 sec, 13 x (98°C for 10 sec, 63°C for 30 sec), 72°C for 1 min). The library PCR products were cleaned up with Agencourt AMPure XP beads (Beckman Coulter, A63881) and sequenced at the Molecular Biology Core Facilities of the Dana-Farber Cancer institute on a NovaSeq sequencer with depths of 10-20 million reads per sample and 150bp Paired-End sequencing.

### RNA analysis

#### RT-qPCR

Total RNA was isolated using the Monarch Total RNA Miniprep Kit (NEB, T2010S). The RT-qPCR assays were performed using 150 ng of total RNA and the Luna Universal One-Step RT-qPCR Kit (NEB, E3005S) on a QuantStudio 3 system (Applied Biosystems). The RT-qPCR conditions were 55°C for 10 min, 95°C for 1 min, 45 cycles of 95°C for 10 sec and 60°C for 1 min. The mRNA amplification was quantified using the comparative CT Method normalized to GAPDH. Primers are listed in **Table S6**.

RNA sequencing. RNA was extracted as above. RNA-seq libraries preparation and sequencing was performed by Genewiz (Azenta) on Illumina® NovaSeq™. Reads were mapped to the hg19 human reference genome, and gene expression levels were quantified using the R package DESeq2 (41).

### Chromatin immunoprecipitation (ChIP) and qPCR

ChIP assays were performed using the SimpleChIP Chromatin IP kit (Cell Signaling Technology). Cells were crosslinked with 1% formaldehyde for 5 min and quenched with 125mM glycine. Cell pellets were solubilized in ChIP buffer (Cell Signaling Technology) and sonicated (Fisher BioRuptor U200). Part of the supernatant was digested with Proteinase K (65°C for 2 hr), the DNA isolated by spin columns, and input DNA quantified by real-time PCR. Equivalent amounts of chromatin were incubated with primary antibody overnight at 4°C followed by protein-G magnetic beads for 2 hr at 4°C (Cell Signaling Technology). Immune complexes were washed in low- and high-salt ChIP buffers (Cell Signaling Technology), eluted and incubated in NaCl (65 °C for 2 hr), and then digested with Proteinase K and purified with the QiaQuick PCR Purification Kit (Qiagen). Purified DNA was quantified by qPCR using Luna Universal qPCR Mastermix (NEB) on the QuantStudio 3 system (Applied Biosystems). ChIP-grade antibodies included RCOR3 (Abcam ab76921) H3K4me2 clone C64G9 (CST#9725), and rabbit IgG control (Abcam ab171870). Primer sequences are listed in **Table S7**.

### Data analysis

#### CUT&Tag

Following a quality control check using FASTQC (v.0.11.9), raw FASTQ reads were aligned to the hg19 and the dm6 reference genomes (downloaded from the UCSC database) using Bowtie2 (v.2.4.5) (42). Mosaic adapter sequences were trimmed during this step using the −5 19 parameters. The SAM files were deduplicated using Picard MarkDuplicates (v.2.27.4) followed by conversion to BAM format using samtools (v.1.15.1) (43) and blacklist removal from the BAM files with bedtools intersect (v.2.30.0) (44) using the ENCODE blacklist bed file for hg19 (45). Genome coverage tracks were generated using deeptools bamCoverage (v.3.5.1) (46) with the following parameters: --binSize 5 --normalizeUsing RPGC --effectiveGenomeSize 2,864,785,220. Peaks were called using MACS2 (v.2.1.4) (46) and high quality peaks were identified as peaks present in two or more replicates generated with bedtools intersect. The BAM files were then converted to bedgraph format using bamCoverage with the same parameters as above and --scaleFactor followed by the calculated scaling factor. Peaks were subset by presence at each timepoint using Intervene (v.0.6.5). Peaks present only at the mock timepoint were denoted “basal” peaks. Any peaks that were present post-UV but not in the mock timepoint were denoted “UV-associated peaks.” For individual post-UV timepoints, bedtools intersect was used to subtract mock timepoint peaks before downstream integration with RNA-seq.

### G4 motif analysis

Peaks were transformed into FASTA files using the bedtools getfasta command, and the sequences were searched for recurrent motifs using MEME suite (47) with the zero or one occurrence per sequence (zoops) option. FASTA sequences were also used to identify G4 motifs using G4Catchall (48) using the following parameters: --G2L 1..12 --max_imperfect_Gtracts 0 --G4H. The absolute values of the G4Hunter scores outputted by G4Catchall were plotted for the different conditions using the ggplot2 R package. The G-run length was calculated for the first four G-runs detected in each motif (5’ to 3’ orientation).

### RNA-seq Differential Expression Analysis

Following a quality control check using FASTQC (v.0.11.9), reads were trimmed with Trimmomatic (v.0.39) and aligned to the indexed Hg19 genome with STAR (v.2.5.2b). Files were sorted and indexed with Samtools (v.1.9) and featureCounts from SubRead (v.2.0.6) was used to count exonic reads per gene. Differential expression analysis was carried out using DESeq2 (1.44.0) where each timepoint post-UV was compared to mock.

### Peak annotation and pathway analysis

UV G4 peaks were annotated using the ChIPseeker R package (v.1.40.0). Gene lists of promoter UV G4 peaks per timepoint were overlapped with lists of corresponding UV DEGs with an adjusted *p*-value (*p*-adj) <0.05. Pathway analysis on those DEGs with promoter UV G4 peaks was done with the ClusterProfiler R package (v4.20.0).

### Proteomic analysis and protein prioritization with ProtFiler

#### Initial mass spectrometry analysis

Mass spectrometry results were pooled per timepoint and deduplicated. Proteins pulled down by miniTurbo control at any timepoint were subtracted from miniTurbo-G4P hits. 502 proteins remained that were found only in miniTurbo-G4P conditions, and these proteins were scored using our protein prioritization tool, ProtFiler. Scoring was based on the following criteria:

#### Relevant Literature

The *Research* function within Entrez Programming Utilities was used to query PubMed in order to gain information on relevant literature. Output included total number of articles based on provided protein symbol (queried as a MeSH term or word within the text), as well as number of articles based on protein symbol and specific search terms included in the title or abstract (“UV”, “ultraviolet radiation”, “quadruplex”, “G4”, “dna repair”, or “melanoma”) to pinpoint relevant, yet novel proteins.

#### String interactions

The STRING database of documented and predicted protein-protein interactions was accessed using STRING API developed by STRINGdb: UniProt identifiers were mapped to STRING identifiers and used to retrieve the interaction network for the given input proteins and gather information about their interconnectedness. The total count of interactions for each protein was calculated in order to prioritize proteins that had greater associations with other proteins in the input set.

#### GO terms

Proteins API (49) was used to query gene ontology terms associated with each protein based on their UniProt identifier. Proteins were weighted based on associated gene ontology terms to capture relevant information about protein function that might be missing or less prevalent in the titles and abstracts searched. RNA-related GO terms received negative weight in order to maintain focus on DNA-protein interactions. Positive and negative terms are listed in **Table S8**. The GO score for each protein was calculated based on the gene ontology terms retrieved from the Proteins API. A GO term was assigned a 1 if it included relevant terms for our use case such as “DNA binding” or “DNA repair”, a -1 if it contained a less desired term like “RNA processing” or “myosin,” and otherwise was assigned 0. These numerical assignments were then summed for every protein.

#### Antibody Availability

To select antibodies for each protein, <u>antibodypedia.com</u> was queried based on given UniProt IDs. Web-scraping and browser automation was carried out with the Selenium package (v.3.141.0) (RRID:SCR_025969, http://www.seleniumhq.org/projects/webdriver); output included antibody link(s) and number of total referenced antibodies.

#### UV Timepoint score

Optional UV score was included to give weight to late-timepoint UV G4 interactors with weights assigned as follows: “mock” as -4, “01h” as 1, “24h” as 2, and “48h” as 3. To calculate the final UV score for each protein, weights for each timepoint were summed.

#### Overall score

Overall score for each protein was calculated based on normalized total article count, search term article count, number of referenced antibodies via Antibodypedia, “String” interactions, GO score, and UV score. Greater positive weight was given to “String” interactions (1.5) and UV score (2x), and negative weight was given to total article count (−1).

### Statistics

Data presented in bar graphs represent the mean ± SD of at least three biological repeats. Statistical analysis was performed using an unpaired two-tailed Student *t* test comparing treatment versus vehicle or one-way ANOVA to compare multiple conditions. A *p*-value of less than 0.05 was considered statistically significant. Immunoblotting results are representative of at least three experiments. Density plots for G4 hunter score and G4 motif length were analyzed using the Wilcoxon test for comparison between two groups of samples. All statistical analyses and visualization were performed with Prism 11 or R (v.4.5.2).

### Data visualization

Genome browser views were created using IGV (v.2.12.3). Scaled Venn diagrams were created using meta-chart (https://www.meta-chart.com/) or eulerr R package (v.7.0.3). Upset plots were created using UpSetR (v.1.4.1). Graphing was done in Prism 11 or using ggplot2 (v.4.0.3).

### Data Availability

The CUT&Tag and RNAseq datasets generated in this study have been deposited in the Zenodo database and files will be made publicly available upon publication (DOI: 10.5281/zenodo.21316659). The hg19 and dm6 reference genomes were retrieved from the UCSC database. The hg19 blacklist file (identifier ENCFF200UUD) was obtained from the ENCODE database.

### Code Availability

Custom scripts developed in Python, R, and Bash for this study, including for ProtFiler, are available at https://github.com/Day-Lab-Bioinformatics. The G4Catchall script was obtained from https://github.com/odoluca/G4Catchall.

## RESULTS

### Persistent G4 DNA Induction Distinguishes UV Among Common Genotoxins

To gain greater insight into the crosstalk between G4 DNA structures and DNA repair processes, we tested a small cohort of varied genotoxins for their ability to influence global G4 DNA formation. We treated A549 lung adenocarcinoma cells with bleomycin to induce DNA double strand breaks (DSBs), methyl methanesulfonate (MMS) to alkylate DNA, camptothecin (CPT) to trap topoisomerase as a protein-DNA adduct, and ultraviolet radiation B (UV-B) to induce lesions including cyclobutane pyrimidine dimers (CPDs). Measuring the levels of G4 DNA in a time course following genotoxin treatment revealed that UV-B was the sole genotoxin that led to a significant increase in global G4 DNA and that the increase persisted at late timepoints (**Fig. 1A, S1**). We observed a similar induction and persistence of UV G4s in human A375 melanoma cells (**Fig. 1B-C**), which are derived from skin, a tissue for which solar UV-B is a ubiquitous and harmful mutagen. CPDs are a major product of UV radiation that activate nucleotide excision repair (NER) to excise the lesions. Interestingly, UV G4s persisted at 48 and 96 hr (**Fig. 1A-C**), exceeding the time needed to complete most NER (50), suggesting that the G4s persist after repair is complete and their persistence may have physiological consequences. Remarkably, we also observed elevated and persistent G4 DNA in a mouse model of moderate sunburn (**Fig 1D-E**) (51), confirming the induction of UV G4s *in vivo*. Taken together, these data suggest that UV-B induces persistent G4 DNA formation across multiple cell types *in vitro* and *in vivo*.

**Figure 1.**
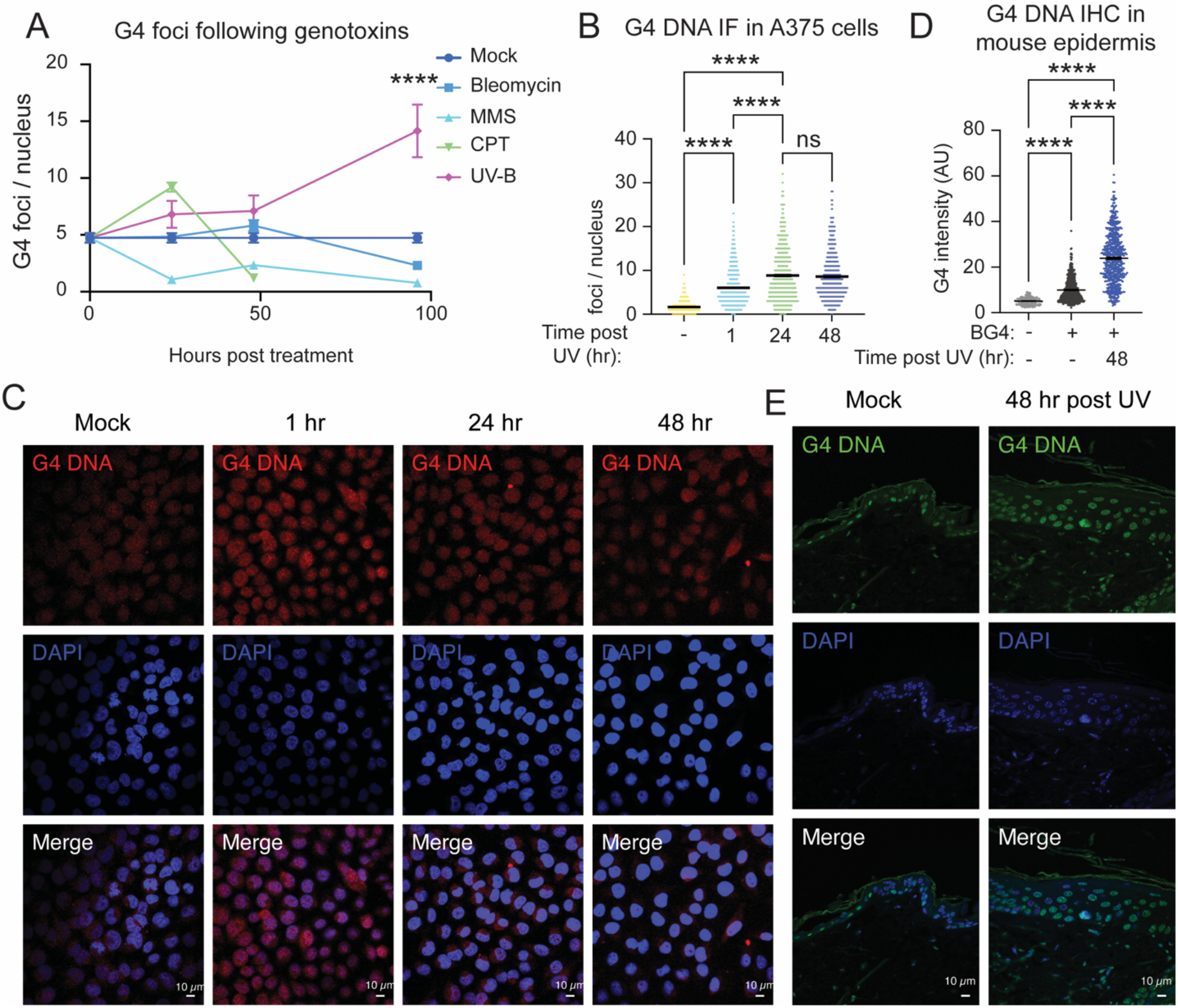
G4 DNA Induction Distinguishes UV Among Common Genotoxins. **A.** Quantitation of BG4 DNA foci by immunofluorescence (IF) following treatment with bleomycin (5 µg/mL), MMS (100 µM), CPT (500 nM), or UV-B (10 J/m^2^) in A549 cells, followed by the indicated recovery times. **B-C.** Quantitation of BG4 DNA foci (**B**) and illustrative images (**C**) of A375 cells following 10J/m^2^ UV-B treatment with the indicated recovery. **D-E.** Quantitation mean immunofluorescence intensity of BG4 (**D**) and illustrative images (**E**) of epidermal cells above the basement membrane in mouse tissue irradiated with 20J/m^2^ with the indicated recovery times. MMS, methyl methanesulfonate; CPT, camptothecin; *, *p* < 0.05; **, *p* < 0.01; ***, *p* < 0.001; ****, *p* < 0.0001 (One-way ANOVA for multiple comparisons).

### UV G4s arise at specific locations throughout the genome

To identify the specific loci whose DNA structures are influenced by UV, we performed G4 CUT&Tag (14,40) at 0, 1, 24 and 48 hr following 10 J/m^2^ UV-B (**Fig. 2A-B**) in A549 cells. We designated high-quality G4 peaks as those found in at least two out of three biological replicates (**Fig. S2A**) and observed that they were enriched for G4-forming motifs (**Fig. S2B**), indicating that our technique allowed us to capture G4 structures. An increase in the total number of high-quality G4 peaks was observed at 1 and 48 hours following UV exposure (**Fig. 2A**), accompanied by a significant increase in G4 peak intensity at 24 and 48 hr (**Fig. 2C**). These findings are consistent with our IF results, which shows elevated G4 signal across all timepoints following UV-B treatment (**Fig. 1A**). We identified all high-quality G4 peaks that emerged following UV exposure (**Fig. 2D**) to compare their characteristics with those present in unirradiated cells. Using G4Hunter, a tool that calculates G4-forming propensity based on G-richness and G-skewness (52), we observed that UV-induced G4s display modest but significantly elevated propensity scores relative to basal G4s (**Fig. 2E**; *p* = 0.00348, Kolmogorov–Smirnov test), suggesting a slight increase in thermodynamic stability. Because shorter loop lengths are associated with greater G4 stability, we next examined overall G4 motif sequence length and found that UV-induced G4s derive from significantly shorter DNA sequences than basal G4s (**Fig. 2F**; *p* = 1.77 × 10⁻¹⁶, Kolmogorov–Smirnov test), indicating that their enhanced stability may be driven by shorter loop regions. Interestingly, UV G4s exhibited a greater number of G repeats (**Fig. S2C**, *p* < 2.67 x 10^-16^, Kolmogorov-Smirnov test), suggesting that a subset of UV G4s may contain a fifth G repeat that has been termed a ‘spare tire’ (53). Taken together, this data suggests that UV G4s emerge at distinct loci throughout the genome and tend towards higher thermodynamic stability than basal G4 structures.

**Figure 2.**
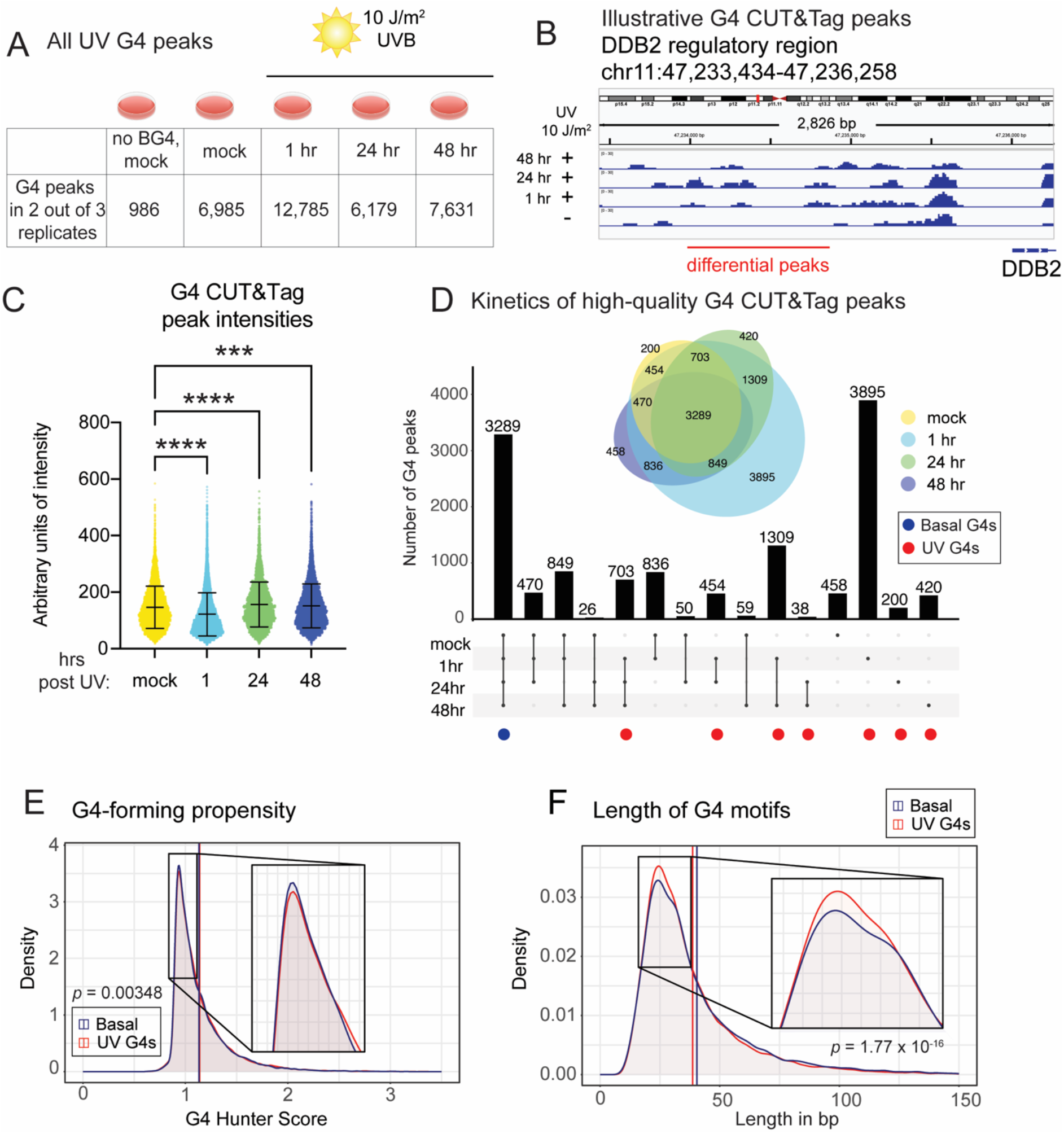
UV G4s have distinct properties and arise at specific locations throughout the genome. **A.** Number of high quality G4 DNA peaks after optional 10 J/m^2^ UV-B followed by indicated recovery times. **B.** Genome browser view of G4 CUT&Tag signal at illustrative locus in the DDB2 promoter region. **C.** Peak intensity values of high quality G4 peaks after optional 10 J/m^2^ UV-B followed by indicated recovery times. **D.** Categorization of overlap of high quality G4 DNA peaks across timepoints. **E.** Density plot of G4-forming propensity based on G4-Hunter score of basal peaks versus UV G4s. **F.** Density plot of G4 motif length based on G4 Catchall analysis of basal peaks versus UV G4 peaks. *p* < 0.05; **, *p* < 0.01; ***, *p* < 0.001; ****, *p* < 0.0001 (One-way ANOVA for multiple comparisons).

### Integration of UV G4 structure landscape with UV-induced transcriptional changes

G4 structures are known to elicit transcriptional influence (17), particularly as positive regulators of transcription (16) when they are located near regulatory regions such as promoters. We performed RNA-seq in A549 cells at 0, 1, 24, and 48 hr following 10 J/m^2^ UV-B (**Fig. 3A-C**, **S3A**) to integrate this data with the profiles of UV G4 peaks described in **Figure 2**. Relative to unirradiated cells (0 hr), we identified 1,220, 613, and 198 differentially expressed genes at 1, 24, and 48 hr with *p*-adjusted value < 0.05 (**Table 1**). Consistent with previous reports (54,55), a larger number of genes were differentially expressed at the earlier timepoint than at later timepoints. Despite cell type-specific differences (55), we noted a handful of well-documented UV-induced genes including PTGS2 and CDKN1A in our RNA-seq datasets (56).

**Figure 3.**
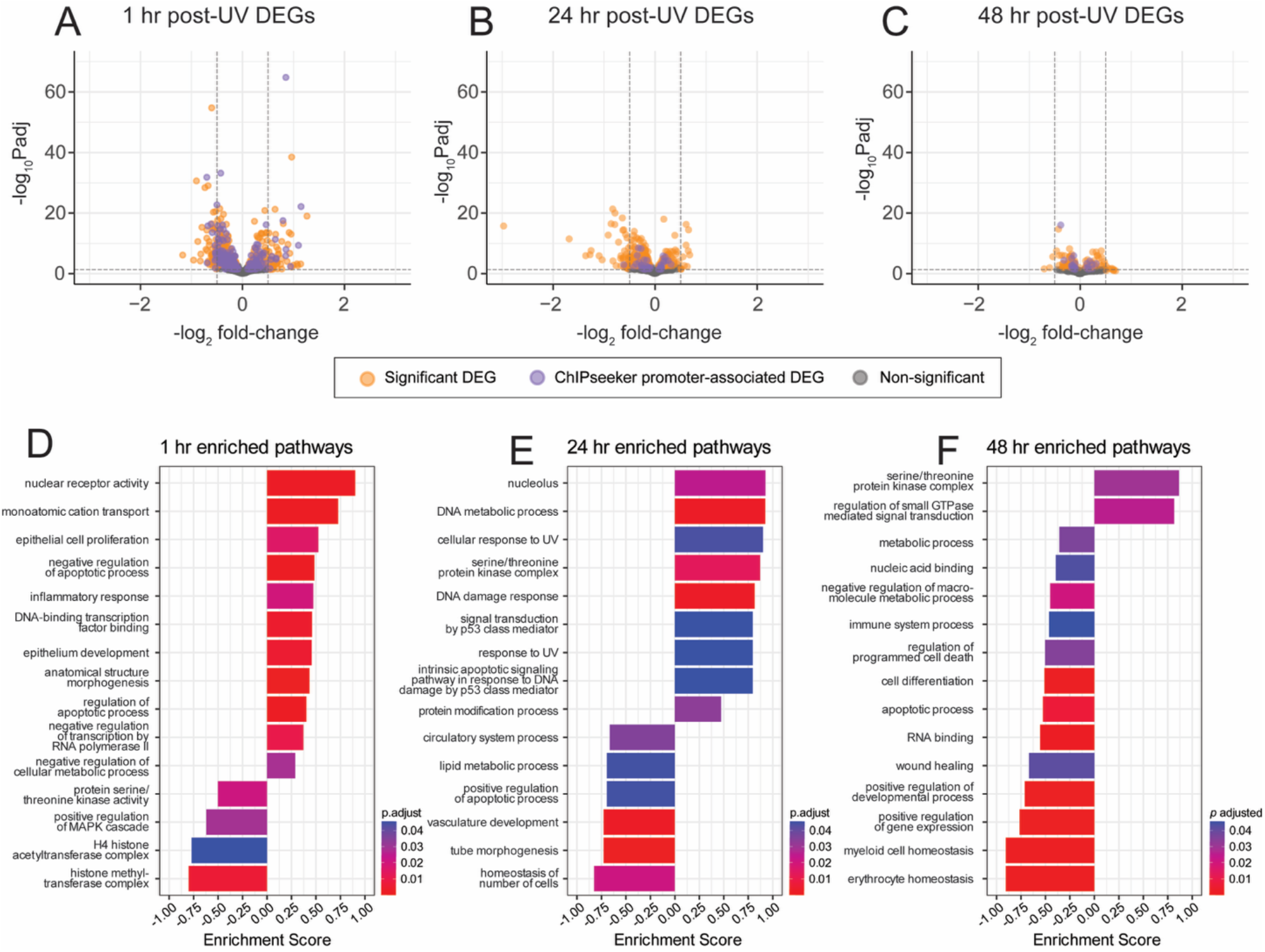
UV G4s influence gene expression and regulate pathways related to damage and repair. A-C. Differentially expressed genes (DEGs) relative to mock following 10 J/m^2^ UV-B at 1 hr (**A**), 24 hr (**B**), and 48 hr (**C**). Promoter-associated genes designated by ChIPSeeker indicated in purple. **D-F.** Enriched pathways identified by Gene Set Enrichment Analysis of Gene Ontology analysis of ChIPSeeker positive genes at 1 hr (**D**), 24 hr (**E**), and 48 hr (**F**). hr, hours.

To identify genes that are likely influenced by UV-induced G4 formation, we used ChIPseeker (57) to map high-quality UV G4 peaks to nearby regulatory elements, including transcription start sites, promoters, and enhancers. We then examined whether these genes showed differential expression at each timepoint post-UV exposure. This approach allowed us to identify 285, 40, and 37 genes, at 1, 24, and 48 hr respectively, whose expression might be influenced by the emergence of a UV G4 in a nearby regulatory sequence (**Table 2**). Gene set enrichment GO analysis (58,59) of the differentially expressed genes that are associated with UV G4s revealed that distinct pathways of interest were enriched at each timepoint (**Fig. 3D-F, S3B-D**).

At 1 hr, genes associated with UV G4s were enriched for apoptotic and inflammatory response pathways, among others (**Fig. 3D**), consistent with the well-established role of UV exposure in triggering apoptosis and inflammation (3,6,60). At 24 hr, significant enrichment was observed for DNA repair and UV response pathways (**Fig. 3E**), including genes such as DDB2, CDKN1A, and BCL3 (**Fig. S3C**). Also at 24 hr, enrichment was seen for vasculature development, in line with prior reports of UV-induced vascular remodeling (4). By 48 hr, UV G4-associated genes were enriched for metabolic processes (**Fig. 3F**), consistent with evidence that UV radiation can dysregulate metabolic function (61). Taken together, this indicates that increased expression of certain genes driven by a UV G4 could contribute to several of the UV-induced phenomena described in the literature. In addition, UV G4s are associated with elevated expression of MELTF (at 48 hr) and AXL (at 24 and 48 hr), two genes that are implicated in the development and progression of melanoma (62,63), suggesting that UV G4s may also drive the expression of genes contributing to UV-induced pathology.

### Integration of UV G4 landscape with the G4 proteome

To address how UV G4s may form and persist, we established a system in human cells to covalently label and characterize the network of proteins spatially proximal to G4s. We fused the peptide G4P (22) to the HA-tagged miniTurbo biotin ligase (33) and placed it under the control of an inducible promoter in A375 human melanoma cells (**Fig. 4A-B**). We noted that the HA-miniTurbo-G4P IF signal was significantly increased in the nucleus at 24 hr following 10 J/m^2^ UV (**Fig. 4C-D**), suggesting that HA-miniTurbo-G4P is associated with UV G4s. Using affinity purification followed by mass spectrometry, we identified proteins associated with G4s at the indicated timepoints following UV-B radiation (**Table S1**). After eliminating all targets that were biotinylated by miniTurbo alone or by both miniTurbo and by miniTurbo-G4P, we identified 502 proteins specifically biotinylated by miniTurbo-G4P (**Fig. 4E**, **Table 3, Fig. S4A**); this allowed us to identify proteins associated with G4s both before and after UV. To prioritize within this large number of proteomically-identified G4-associated factors, we developed ProtFiler, a flexible computational tool that integrates publicly available data to assign a prioritization score (**Fig. S4B**). ProtFiler gathers information on protein network connectivity strength, reagent availability, GO terms, and relevant keywords from resources such as STRING (64), PubMed, EMBL-EBI, and Antibodypedia (65). In our analysis, proteins are assigned higher scores if they 1) appeared at late timepoints post UV, 2) strongly interacted with other identified hits, 3) were linked to desirable GO terms (e.g. DNA repair) or key words (e.g. UV radiation) and 4) had readily available antibodies. In contrast, proteins that were detected even in the absence of UV or associated with undesirable GO terms or key words (e.g. RNA processing) received lower scores. To focus on less-studied proteins, those with substantially high numbers of corresponding PubMed abstracts were also deprioritized.

**Figure 4.**
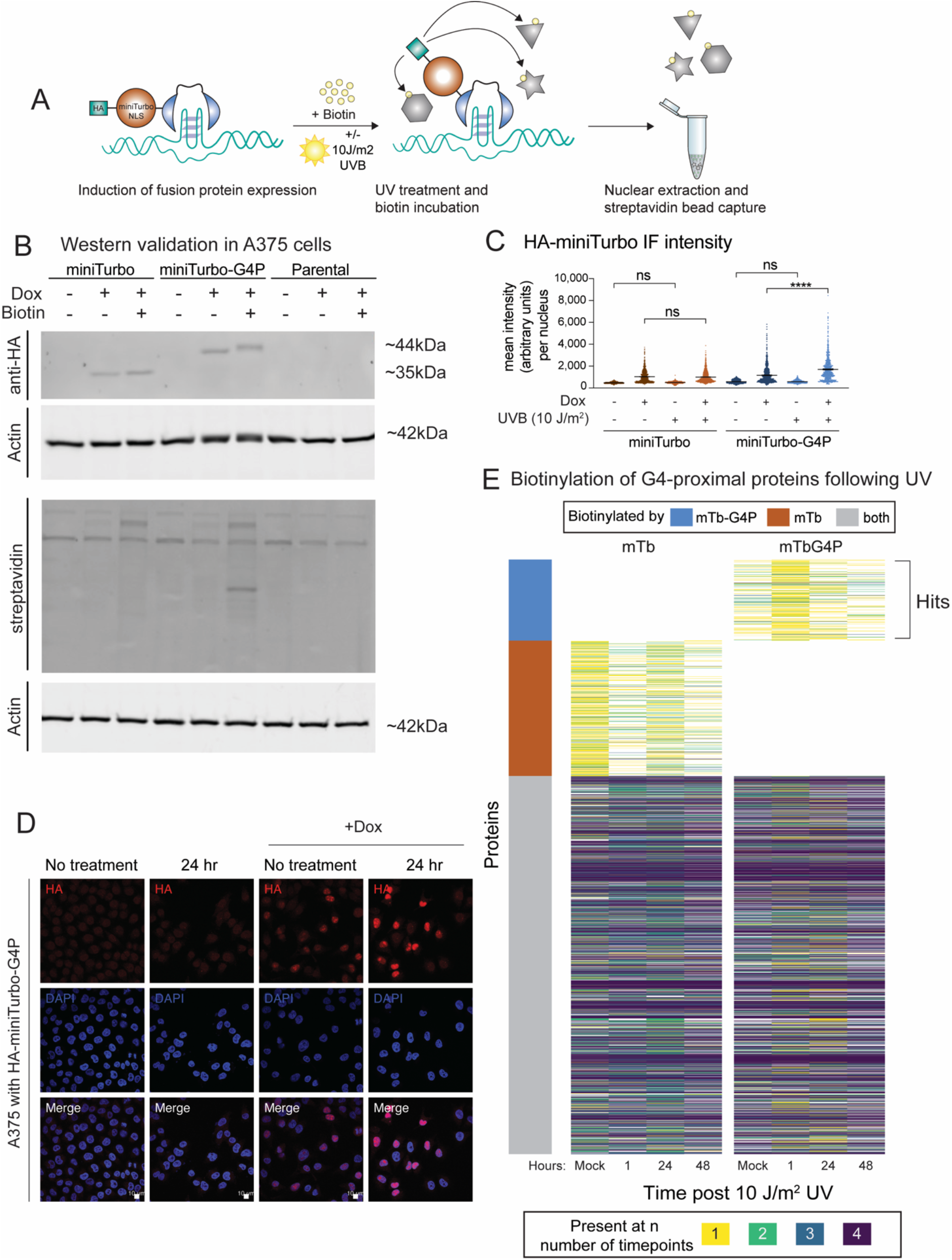
Identification of the UV-induced G4 interactome. **A.** Schematic representation of HA-tagged mTb-G4P expression, biotinylation, and streptavidin protein pulldown. **B.** Western blot of mTb-G4P or mTb expression in A549 cells. Cells were optionally treated with doxycycline (2 µg/mL) to induce fusion protein expression and incubated with biotin for 30 minutes as indicated to allow biotinylation of proximal proteins. anti-HA shows HA-mTb-G4P fusion protein expression and anti-streptavidin shows biotinylation of nuclear extracted protein. **C-D.** Quantitation (**C**) and illustrative images (**D**) of HA-miniTurbo or HA-miniTurbo-G4P IF mean intensity with doxycycline for 24 hr and 10 J/m^2^ UV-B treatment as indicated. **E.** Heatmap of proteins identified by mass spectrometry in mTb-G4P and mTb alone conditions after optional treatment with 10 J/m^2^ UV-B and indicated recovery times. Blue bar on the left indicates the 502 proteins present exclusively in mTb-G4P, orange bar indicates those only present only in the mTb, and gray indicates those present in both. mTb, miniTurbo; mTb-G4P, miniTurbo-G4P fusion. ns, not significant. *, *p* < 0.05; **, *p* < 0.01; ***, *p* < 0.001; ****, *p* < 0.0001 (One-way ANOVA for multiple comparisons).

Using Protfiler, we generated a rank-ordered list of the 502 proteins identified by miniTurbo-G4P (**Fig. S4C, Table 4**) and selected 13 of the top-scoring candidates (**Fig. 5A-B**) to assess their functional roles in UV-induced G4 formation. ZRF1, which was identified proteomically in our screen at 24 hr, has been reported to promote short-term (8 hr) induction of UV G4s (8,9) and was therefore included as a positive control. We depleted each of the 13 candidates by siRNA in A375 melanoma cells and found that, consistent with expectations, ZRF1 depletion suppressed UV G4 formation (**Fig. 5B-F**). Notably, depletion of 10 of the 13 newly identified G4-associated factors similarly prevented UV G4 formation, to varying degrees (**Fig. 5B**). We also note that in all cases, depletion of G4-factors led to basal increases in G4 formation (**Fig. S5A**), suggesting that each of these factors also plays a role in regulating G4 formation in the absence of UV exposure. Collectively, these results define a set of genetic dependencies underlying UV G4 formation, prompting us to focus subsequent mechanistic studies on one of the factors in this group, RCOR3, whose depletion led to a dramatic suppression of UV G4s.

**Figure 5.**
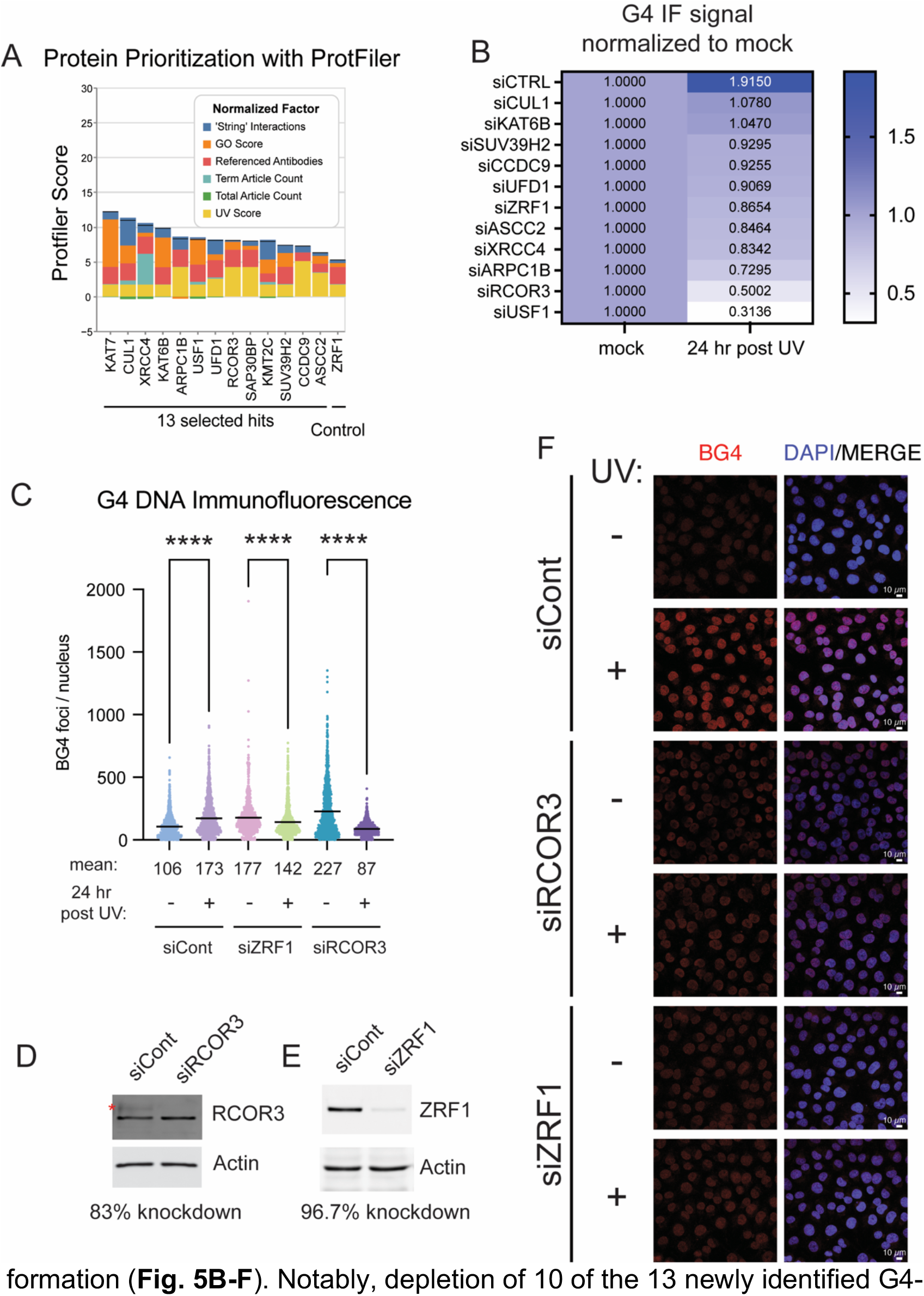
RCOR3 promotes formation of UV G4s. **A.** Composition of ProtFiler scores of the top-ranked proteins that were selected for further analysis. **B.** Heatmap indicating the normalized intensities of BG4 immunofluorescence upon depletion of each of the indicated factors and 24 hr after treatment with 10 J/m^2^ UV-B. Values are normalized to siControl. **C.** Quantitation of BG4 foci / nucleus in cells treated with siControl, siZRF1, and siRCOR3, with 10 J/m^2^ UV-B where indicated. **D-E.** Western blot of siRNA-mediated depletion of RCOR3 (**D**) and ZRF1 (**E**) in A375 cells. **F.** Illustrative BG4 IF images corresponding to the conditions in **C**. *, *p* < 0.05; **, *p* < 0.01; ***, *p* < 0.001; ****, *p* < 0.0001 (One-way ANOVA for multiple comparisons).

### RCOR3 promotes both formation of UV G4s and repair of UV damage

We selected RCOR3 for further study because, as a component of the CoREST chromatin-modifying complex, it represents a plausible regulatory mechanism linking chromatin state to G4 DNA dynamics. Notably, RCOR3 depletion increased basal G4 levels in the absence of UV (**Fig. S5A**) yet produced one of the most pronounced reductions in UV-induced G4s among all the candidate factors (**Fig. 5B**). Not only were UV G4s prevented from forming in the absence of RCOR3, but G4 levels were significantly reduced below baseline (**Fig. 5C-D**). In order to examine the behavior of specific genes, we focused on two genes, DDB2 and CDKN1A/p21, that had both UV-induced G4s in their regulatory regions (**Fig. 6A,B**) and exhibited increased expression at 24 hr post UV. Both DDB2 and CDKN1A/p21 are known targets of p53 (66,67), but we noted that RCOR3 depletion by CRISPRi (68) did not lead to a reduction in p53 either before or after UV (**Fig. S5B-C**), indicating that RCOR3-dependent stabilization of UV G4s is p53-independent.

**Figure 6.**
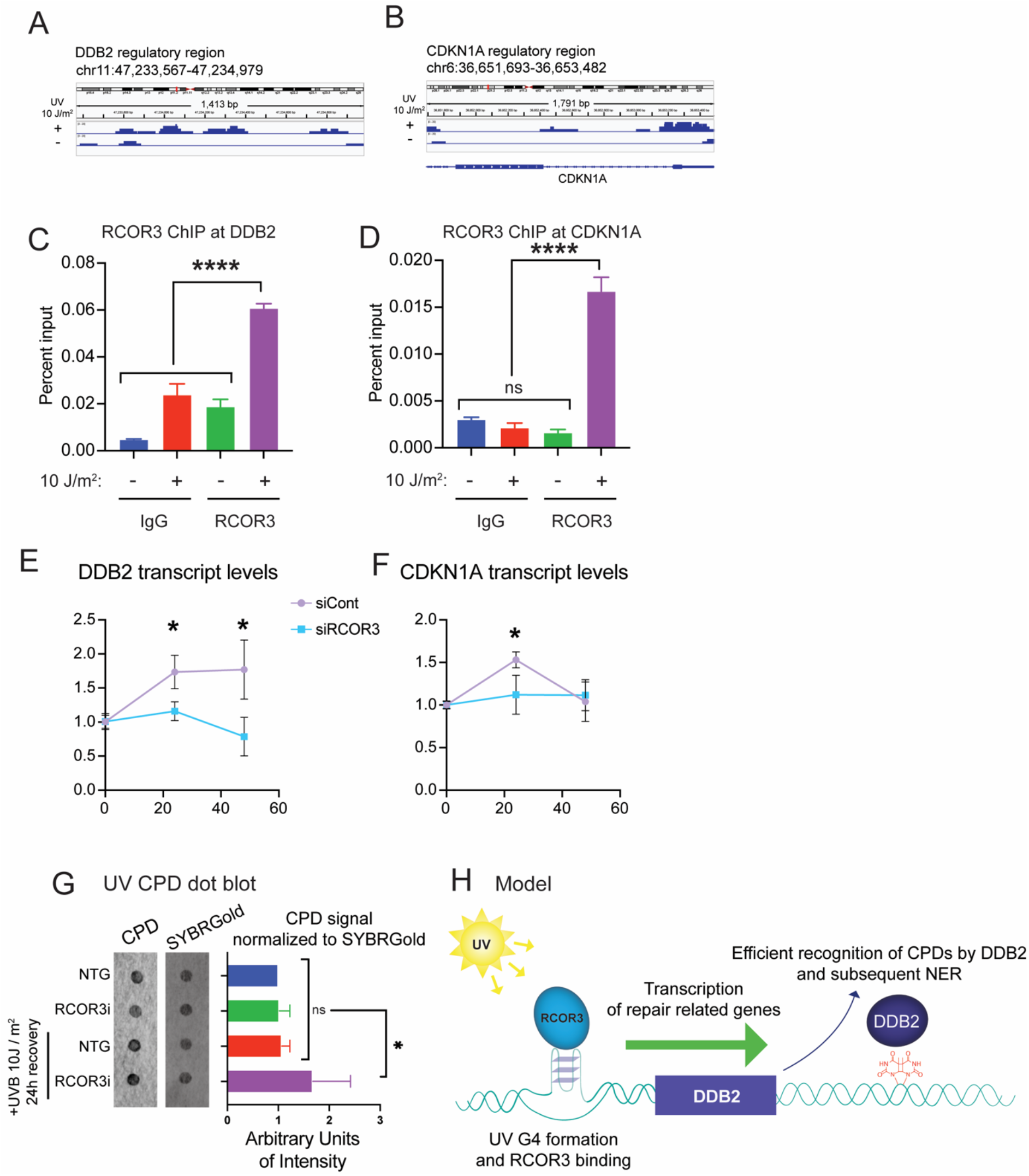
RCOR3 binds and promotes expression of genes associated with UV G4s loci to promote repair of UV damage. **A-B.** Genome browser view of G4 peaks at 0 (-) and 24 hr (+) following 10 J/m^2^ UV at the DDB2 (**A**) and CDKN1A (**B**) regulatory regions. **C-D.** Quantitation of RCOR3 occupancy by ChIP at DDB2 (**C**) and CDKN1A (**D**) G4 peaks in their regulatory regions. **E-F.** Quantitation of DDB2 (**E**) and CDKN1A (**F**) transcript levels after siRCOR3 treatment, compared to siControl in A375 cells. **G.** CPD dot blot and quantitation of CPD signal normalized to SYBR Gold signal after CRISPRi of RCOR3 in A375 cells after optional treatment with 10 J/m^2^ UV-B. **H.** UV-B irradiation triggers RCOR3 binding at specific promoter loci, promoting formation of UV-induced G-quadruplexes (UV G4s) at genes involved in the DNA damage response. G4 formation within these promoters drives expression of target genes, including DDB2, thereby supporting efficient repair of UV-induced lesions. ChIP, chromatin immunoprecipitation. CPD, cyclobutane pyrimidine dimer.

Chromatin immunoprecipitation (ChIP) revealed that 24 hr after UV, RCOR3 exhibited significant recruitment to the UV G4 loci associated with DDB2 and CDKN1A (**Fig. 6C,D**) but not to an unrelated genomic locus (**Fig. S5D**). We subsequently measured the UV-dependent gene expression and found that both DDB2 and CDKN1A exhibited significantly increased expression after UV irradiation as expected but that this increase was abrogated by RCOR3 depletion (**Fig. 6E,F**) with a similar effect observed in A549 cells (**Fig. S6A-C**). Similar findings at loci identified using less stringent G4 CUT&Tag analysis parameters (**Fig. S6D-F**) suggest that the role of UV-induced, RCOR3-supported G4s in regulating gene transcription is robust, and that our primary analysis likely underestimates the full scope of this mechanism.

While RCOR1 activates lysine-specific demethylase (LSD1) to remove methyl groups from H3K4-dimethylated histones (69,70), a biochemical study of the RCOR proteins suggests that RCOR3, in contrast to RCOR1, represses LSD1 activity (71). To test this observation at sites of UV-dependent RCOR3 recruitment in cells, we examined ChIP signals for H3K4-methylation, the primary target of LSD1. Consistent with the biochemical evidence (71), sites of RCOR3 recruitment did not exhibit significant changes in H3K4me2 (**Fig. S7A-C**). We reasoned that UV G4-dependent up-regulation of UV response and DNA repair genes (**Fig. 3E**) might enhance repair of UV lesions. In particular, both DDB2 and CDKN1A play important roles in NER (72–74). In that case, the failure to up-regulate UV repair genes upon RCOR3 depletion would lead to less efficient repair of UV lesions such as CPDs. Indeed, CPDs were elevated and persistent at 24 hr post UV in RCOR3-depleted cells relative to control (**Fig. 6G**). Taken together, these data suggest that RCOR3 stabilization of G4s makes an important contribution to efficient repair of UV lesions.

## DISCUSSION

In this study, we identify UV as a distinct genotoxic stressor capable of inducing G4 DNA structures, a property not shared by the other genotoxins tested. Extending this phenomenon beyond its original characterization (8,9), we demonstrate UV-induced G4 formation across multiple transformed human cell lines and, notably, in mouse epidermis *in vivo*. Interestingly, we find that these G4 structures are not transient but that they persist at late timepoints following UV exposure. Integration of the genome-wide landscape of UV G4s with UV-induced transcriptional alterations and the UV-induced G4 proteome allows us to investigate the mechanistic basis and functional consequences of this persistent UV G4 landscape. After demonstrating that several of the newly identified G4-proximal proteins are important for the formation of UV G4s, we focus on RCOR3 as a key factor that promotes UV G4 formation. Our data indicate that the behavior of RCOR3 on chromatin is fundamentally different from its paralog RCOR1, a core component of a well-studied transcriptional repressor complex. Instead, our data show that UV-induced recruitment of RCOR3 supports elevated gene expression at genes containing UV G4s. Many of the genes associated with UV G4s play roles in DNA repair, and ultimately, we find that RCOR3-dependent gene expression following UV is important for efficient repair of UV-induced DNA lesions. Together, these data suggest that UV G4s promote gene expression that supports efficient repair of UV lesions (**Fig. 6H**).

The finding that RCOR3 promotes gene expression was unexpected, given that RCOR3 belongs to the CoREST family of proteins, which have been studied in the context of transcriptional repression. However, we found that the presence of RCOR3 is important for transcriptional up-regulation following UV (**Fig. 6E,F**). The differences between the activities of RCOR1 and RCOR3 can be understood by comparing the structure of their domains. RCOR1 has both a conserved linker domain through which it interacts with LSD1 and a SANT2 domain which activates LSD1 to remove dimethyl groups from histone H3 lysine 4 (H3K4) (69,70); this demethylation contributes to transcriptional repression. While RCOR3 has a conserved linker domain and can associate with LSD1 (71), it lacks both the SANT2 domain and the capacity to facilitate LSD1-mediated nucleosomal demethylation (71). We have shown that RCOR3, unlike RCOR1, does not activate LSD1, which is consistent with *in vitro* findings (71). Instead, RCOR3 recruitment behaves fundamentally differently from the repressive CoREST complex defined by RCOR1. How RCOR3 supports elevated gene expression remains an open question: it may bind and stabilize the G4 structure itself, recruit a G4-stabilizing protein, or recruit chromatin-modifying enzymes that create a locally permissive transcriptional environment. Future studies will distinguish among these possibilities.

Beyond RCOR3, damage occurring directly within G-rich sequences could offer another mechanism driving UV G4 formation. One possibility is that CPDs with an *anti* structure (i.e. head-to-tail thymines) could form between thymines on adjacent G4 loops; several recent studies have provided evidence that this intriguing phenomenon is chemically possible (75,76), suggesting that UV G4s might be stabilized by anti-CPDs that form on non-adjacent thymines. Alternatively, UV-B also leads to ROS that produce 8-oxoguanine (77), a DNA modification favored in guanine-rich sequences owing to their low redox potential (29). Although the effect of 8-oxoguanine on G4 stability is unresolved (78–81), the presence of these modifications in a G4 motif can stabilize G4 structures in some cases (82,83). Therefore, the presence of 8-oxoguanine modifications in UV G4s could possibly contribute to formation and persistence of UV G4s. Finally, the conversion of 8-oxoguanine to an abasic site by the base excision repair pathway has been proposed as a switch that leads to the formation of a G4 structure (84) and supports elevated gene expression which might be consistent with our findings as well. The possible involvement of DNA damage directly in the G-rich sequence that is stabilized as a UV G4 merits exploration in future experiments.

A recent study in *S. cerevisiae* found that UV induces G4 DNA and that a factor called Zuo1 binds to G4s and stabilizes their formation (8). A follow-up study found that ZRF1, the human homolog of Zuo1, is recruited to UV-induced G4s (9). Our proteomic data supports and extends these findings by identifying ZRF1 in close proximity to UV G4s at 24 hours post-UV (**Fig. S4**), a late timepoint at which ZRF1 is also required for UV G4 persistence (**Fig. 5B**). In yeast, UV G4s are important for the recruitment of NER factors (8). Although we did not detect NER factors besides ZRF1 in our proteomic dataset, this absence is not necessarily informative, as our experimental approach can only confirm the presence of a protein and not its absence. Notably, both the yeast study (8) and our data provide evidence that UV G4s promote NER, albeit by different mechanisms: recruitment of NER factors to chromatin (their study) and up-regulation of DNA repair genes (our study). We note that these mechanisms are not mutually exclusive. Interestingly, budding yeast cells that have been previously exposed to a low dosage of UV radiation exhibit dramatically increased survival following subsequent UV exposure (85). The mechanism underlying this increased UV resistance remains unclear, though histone modifications or other epigenetic marks have been proposed as candidates. One intriguing possibility is that UV G4s contribute to this ongoing resistance, a hypothesis is plausible given the apparent conservation of UV G4 formation from yeast to mice to human.

Taken together, our data support a model in which low-dose UV radiation induces widespread, persistent G4 DNA formation across the genome, dependent on RCOR3 and other factors. These UV G4s form in regulatory regions of many genes, several of which are UV-response genes, including DDB2, driving their increased expression at 24 hours post-UV. RCOR3 depletion abrogates UV G4 formation, blocks this induction of gene expression, and impairs the repair of UV lesions. Together, these data indicate that UV-induced RCOR3 recruitment stabilizes G4s at specific loci to promote expression of genes needed for efficient repair.

## ACKNOWLEDGEMENTS

We thank Elizabeth Crane and Justin Crane for guidance and support in the mouse IHC experiments. We value the feedback and thoughtful insights from all members of the Day lab at Northeastern.

## AUTHOR CONTRIBUTIONS

Lindsay R. Julio: Conceptualization, Data curation, Formal analysis, Investigation, Project administration, Visualization, Writing – review & editing. Diana V. Turrieta Vejar: Investigation. Charlotte H. Walborsky: Investigation. Annika Salpukas: Data curation, Formal analysis, Methodology, Visualization. Ella Chee: Data curation, Formal analysis, Methodology, Visualization. Julianne C. Murthy: Formal analysis, Visualization. Shannon G. MacLeod: Investigation. Adrianna L. Vandeuren: Investigation. Mynaja Ferguson: Data curation, Formal analysis, Investigation. Rachel E. Muriph: Data curation, Formal analysis, Investigation. Jason J. Evans: Data curation, Formal analysis, Investigation. Tovah A. Day: Conceptualization, Funding acquisition, Investigation, Project administration, Supervision, Writing – original draft, Writing – review & editing

## SUPPLEMENTARY DATA

Supplementary Data are available at NAR online.

## DATA AVAILABILITY

The data underlying this article are available in Zenodo at [DOI: 10.5281/zenodo.21316659]. All other data generated during this study are included in this published article and its supplementary information files.

