## Supplementary Figures for "G4 DNA structures induced by UV radiation: a multi-omic approach"

**Supplementary Figures 1-7**

**Supplementary Tables 1-8**

### Figure S1

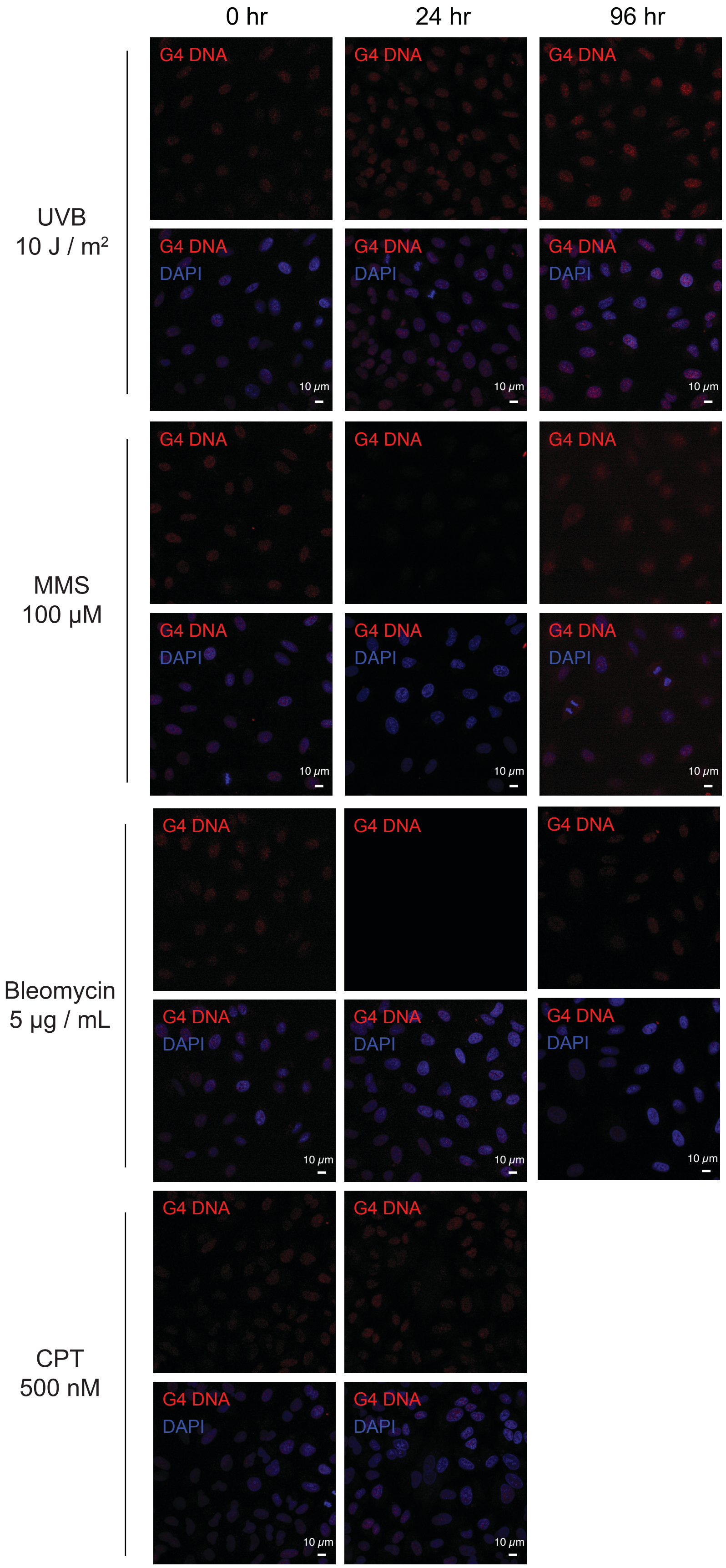

0 hr
24 hr
96 hr
UVB
10 J / m²
MMS
100 μM
Bleomycin
5 μg / mL
CPT
500 nM
G4 DNA
DAPI
10 μm

**Supplementary Figure 1**. **BG4 immunofluorescence examples for genotoxin treatments.** Illustrative BG4 immunofluorescence with RNase treatment of A549 cells at the indicated timepoint following treatment with each genotoxin. MMS, methyl methanesulfonate. CPT, camptothecin.

### Figure S2

A

G4 CUT&Tag replicates

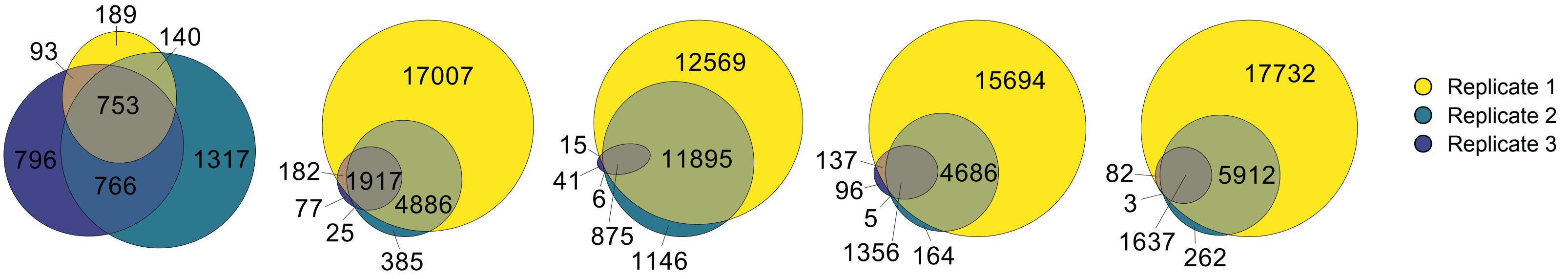


B

| Condition | MEME Suite motifs | E value | Occurrences |
|---|---|---|---|
| 1 hr | 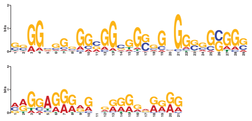 | 7.7e-042<br>2.5e-040 | 7031<br>9960 |
| 24 hr | 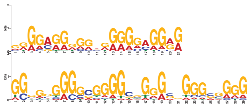 | 2.8e-048<br>7.2e-024 | 5486<br>2957 |
| 48 hr | 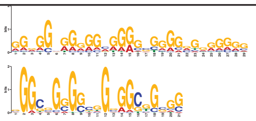 | 1.3e-055<br>6.3e-023 | 6494<br>3908 |
| mock | 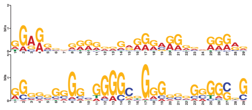 | 2.9e-044<br>3.1e-080 | 4654<br>6528 |
| mock no BG4 | 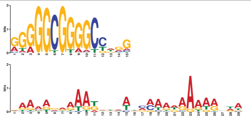 | 2.0e-068<br>1.7e-032 | 460<br>133 |

C

G repeats per G4 motif

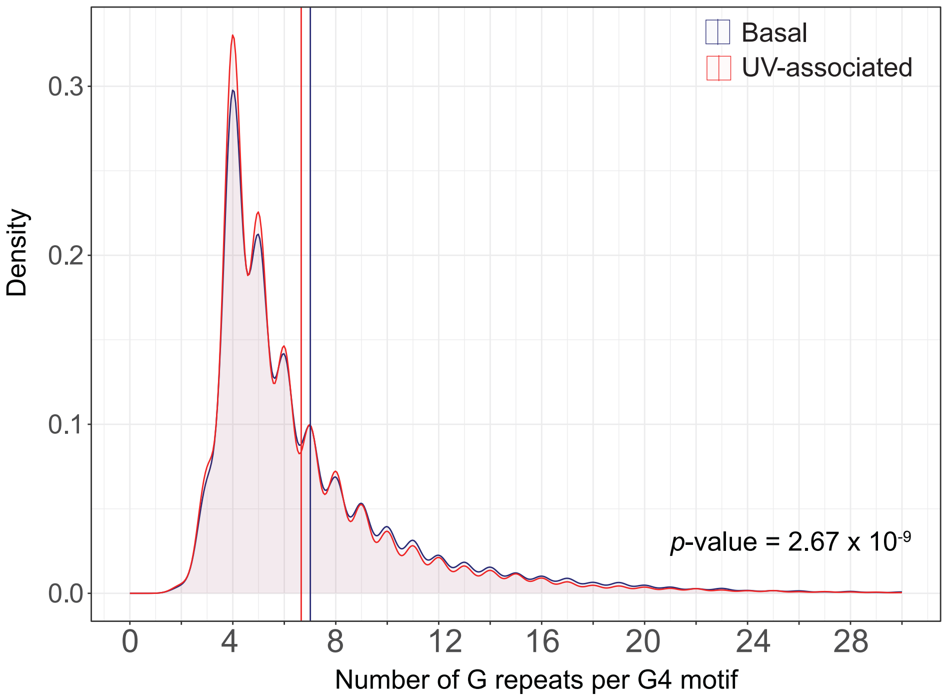

**Supplementary Figure 2. Determination and characterization of high quality G4 CUT&Tag peaks.** A. Venn diagrams representing the three biological replicates used to generate high-quality G4 CUT&Tag peaks, defined as those found in two out of three replicates. Total peaks for each replicate are noted. B. The top two MEME Suite motifs identified in each condition. C. Density plot representing the proportion of G4 motifs found in G4 peaks with the indicated number of repeats.

### Figure S3

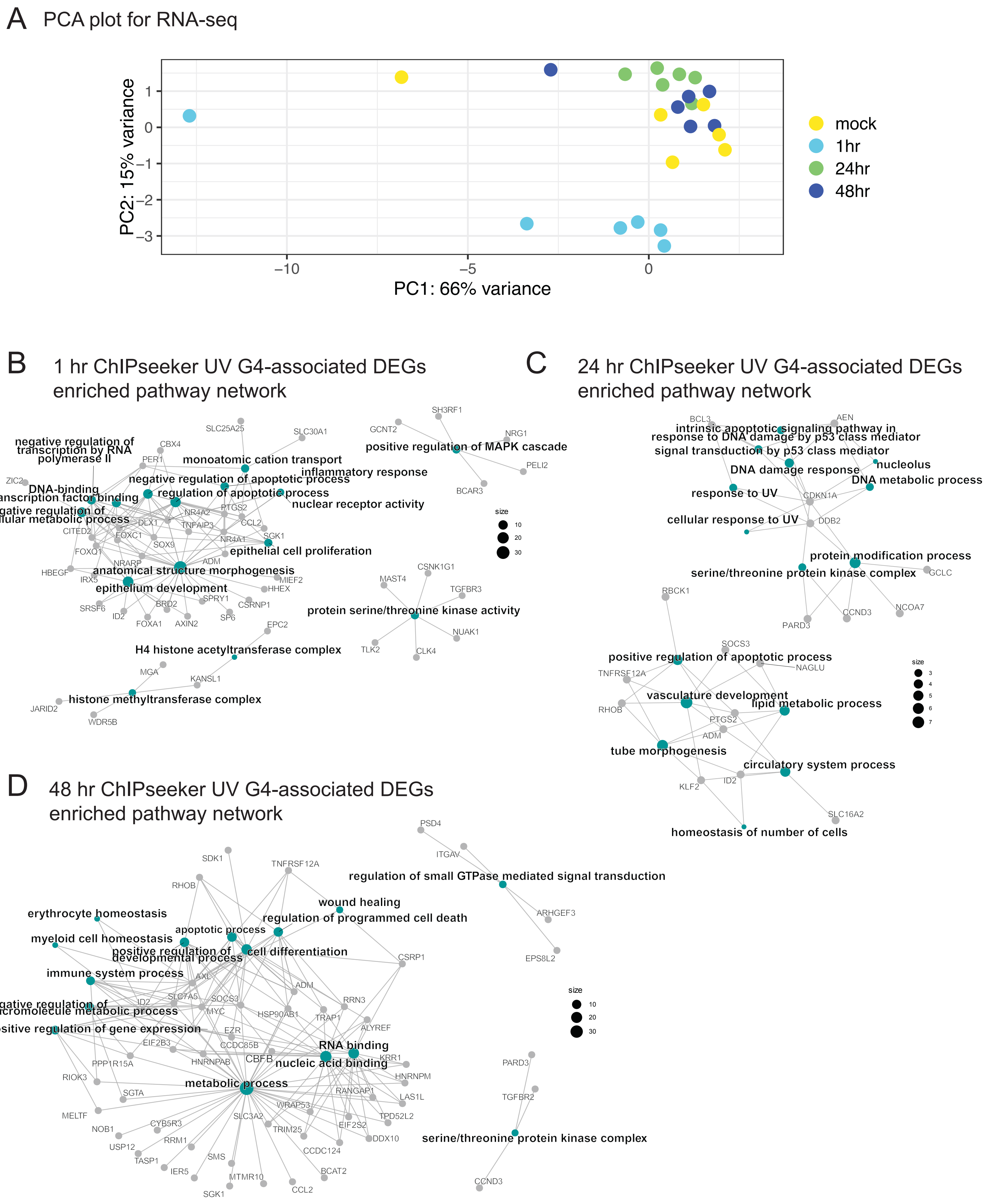

A
PCA plot for RNA-seq
PC2: 15% variance
PC1: 66% variance
1
0
−1
−2
−3
−10
−5
0
mock
1hr
24hr
48hr
B
1 hr ChIPseeker UV G4-associated DEGs enriched pathway network
negative regulation of transcription by RNA polymerase II
monoatomic cation transport
negative regulation of apoptotic process
inflammatory response
DNA-binding transcription factor binding
regulation of apoptotic process
nuclear receptor activity
negative regulation of cellular metabolic process
epithelial cell proliferation
anatomical structure morphogenesis
epithelium development
positive regulation of MAPK cascade
protein serine/threonine kinase activity
H4 histone acetyltransferase complex
histone methyltransferase complex
SLC25A25
SLC30A1
CBX4
PER1
ZIC2
NR4A2
PTGS2
DLX1
TNFAIP3
CCL2
CITED2
FOXC1
NR4A1
SGK1
SOX9
FOXQ1
NRARP
ADM
HBEGF
IRX5
MIEF2
HHEX
SPRY1
CSRNP1
SRSF6
BRD2
ID2
SP6
FOXA1
AXIN2
GCNT2
SH3RF1
NRG1
PELI2
BCAR3
MAST4
CSNK1G1
TGFBR3
NUAK1
TLK2
CLK4
EPC2
MGA
KANSL1
JARID2
WDR5B
size
10
20
30
C
24 hr ChIPseeker UV G4-associated DEGs enriched pathway network
intrinsic apoptotic signaling pathway in response to DNA damage by p53 class mediator
signal transduction by p53 class mediator
nucleolus
DNA damage response
DNA metabolic process
response to UV
cellular response to UV
protein modification process
serine/threonine protein kinase complex
positive regulation of apoptotic process
vasculature development
lipid metabolic process
tube morphogenesis
circulatory system process
homeostasis of number of cells
BCL3
AEN
CDKN1A
DDB2
GCLC
RBCK1
NCOA7
CCND3
PARD3
SOCS3
NAGLU
TNFRSF12A
RHOB
PTGS2
ADM
KLF2
ID2
SLC16A2
size
3
4
5
6
7
D
48 hr ChIPseeker UV G4-associated DEGs enriched pathway network
regulation of small GTPase mediated signal transduction
wound healing
erythrocyte homeostasis
regulation of programmed cell death
apoptotic process
myeloid cell homeostasis
positive regulation of developmental process
cell differentiation
immune system process
negative regulation of macromolecule metabolic process
positive regulation of gene expression
RNA binding
nucleic acid binding
metabolic process
serine/threonine protein kinase complex
PSD4
ITGAV
ARHGEF3
EPS8L2
SDK1
TNFRSF12A
RHOB
CSRP1
AXL
ADM
ID2
SLC7A5
SOCS3
MYC
HSP90AB1
RRN3
TRAP1
ALYREF
EZR
EIF2B3
CCDC85B
CBFB
HNRNPAB
KRR1
PPP1R15A
HNRNPM
RIOK3
SGTA
RANGAP1
LAS1L
MELTF
WRAP53
SLC3A2
EIF2S2
TPD52L2
CYB5R3
NOB1
TRIM25
DDX10
RRM1
USP12
CCDC124
TASP1
SMS
IER5
BCAT2
MTMR10
SGK1
CCL2
PARD3
TGFBR2
CCND3
size
10
20
30

**Supplementary Figure 3. Characterization of transcriptomic profiling.** A. PCA analysis of RNA-sequencing replicates. B-D. Gene-Concept Network Plots for UV G4-associated genes with differential expression at 1 hr (B), 24 hr (C), and 48 hr (D). PCA, principal component analysis. DEGs, differentially expressed genes.

### Figure S4

A

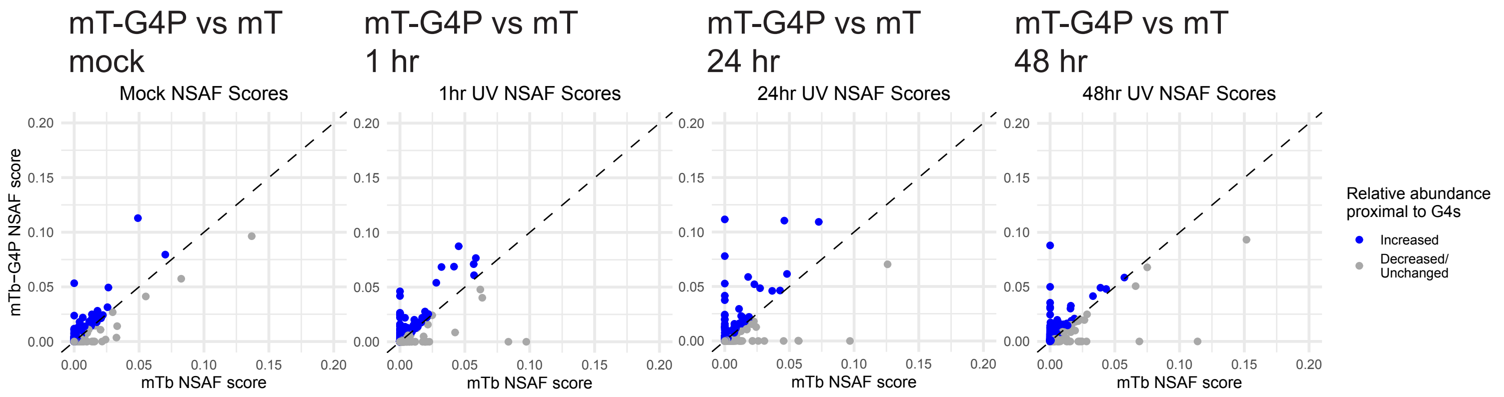


B Schematic of ProtFiler workflow

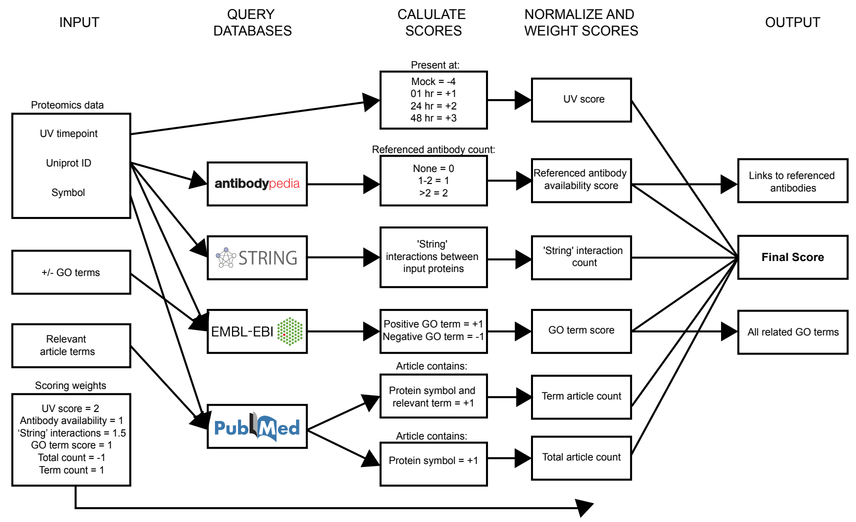


C

ProtFiler scores of all G4-associated proteins

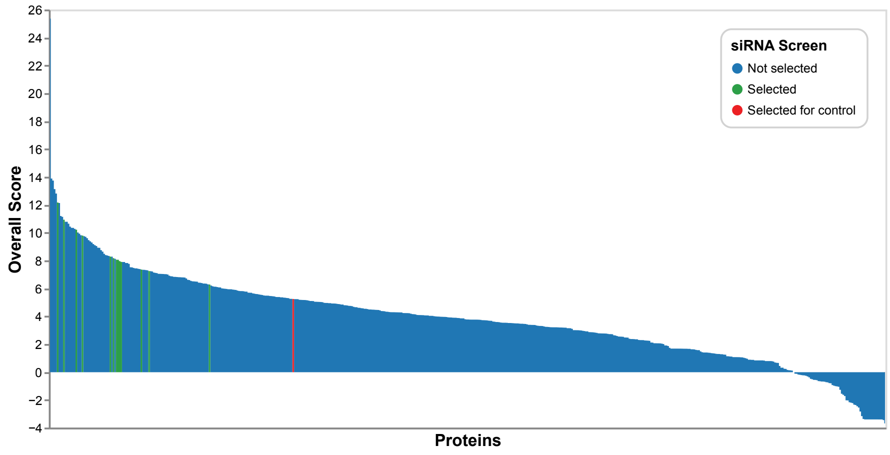

**Supplementary Figure 4. Analysis of G4P-miniTurbo proteomic data.** A. Scatterplots of mT-G4P versus mT NSAF scores for each factor at the indicated timepoints. B. Schematic workflow of the informatic tool ProtFiler that was used to prioritize the list of 514 proteins identified by MS. C. Waterfall plot of ProtFiler scores for each proteomically identified protein. Higher score indicates stronger prioritization to study. mT, miniTurbo. NSAF, normalized spectral abundance factor. MS, mass spectrometry.

### Figure S5

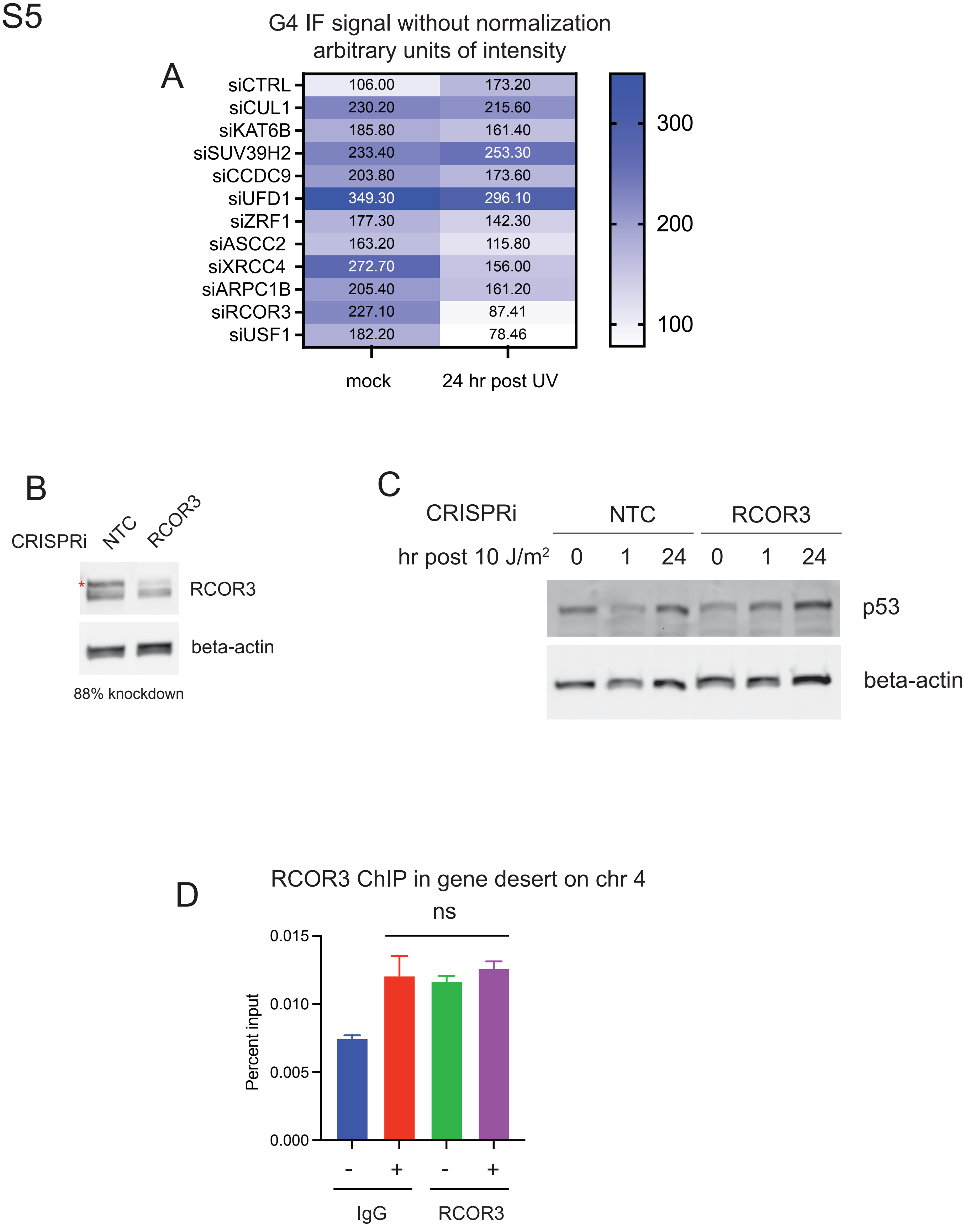
A
G4 IF signal without normalization
arbitrary units of intensity
siCTRL 106.00 173.20
siCUL1 230.20 215.60
siKAT6B 185.80 161.40
siSUV39H2 233.40 253.30
siCCDC9 203.80 173.60
siUFD1 349.30 296.10
siZRF1 177.30 142.30
siASCC2 163.20 115.80
siXRCC4 272.70 156.00
siARPC1B 205.40 161.20
siRCOR3 227.10 87.41
siUSF1 182.20 78.46
mock
24 hr post UV
300
200
100
B
CRISPRi
NTC
RCOR3
*
RCOR3
beta-actin
88% knockdown
C
CRISPRi
NTC
RCOR3
hr post 10 J/m2
0 1 24 0 1 24
p53
beta-actin
D
RCOR3 ChIP in gene desert on chr 4
ns
Percent input
0.015
0.010
0.005
0.000
- + - +
IgG
RCOR3

**Supplementary Figure 5. Characterization of genetic dependence of UV G4 formation.** A. Heatmap indicating the raw intensities of BG4 immunofluorescence upon depletion of each of the indicated factors and treatment with 10 J/m$^2$ UV. Scale is in arbitrary units of intensity. B. Western blot of RCOR3 in A375 cells expressing CRISPRi with either NTC or RCOR3. Asterisk shows the RCOR3 band and percent knockdown is calculated by normalization to beta-actin. C. Western blot of p53 with optional CRISPRi-mediated depletion of RCOR3 following UV at the indicated timepoints. D. RCOR3 occupancy by ChIP at a gene desert on chromosome 4. NTC, non-targeting control. ChIP, chromatin immunoprecipitation. ns, not significant.

### Figure S6

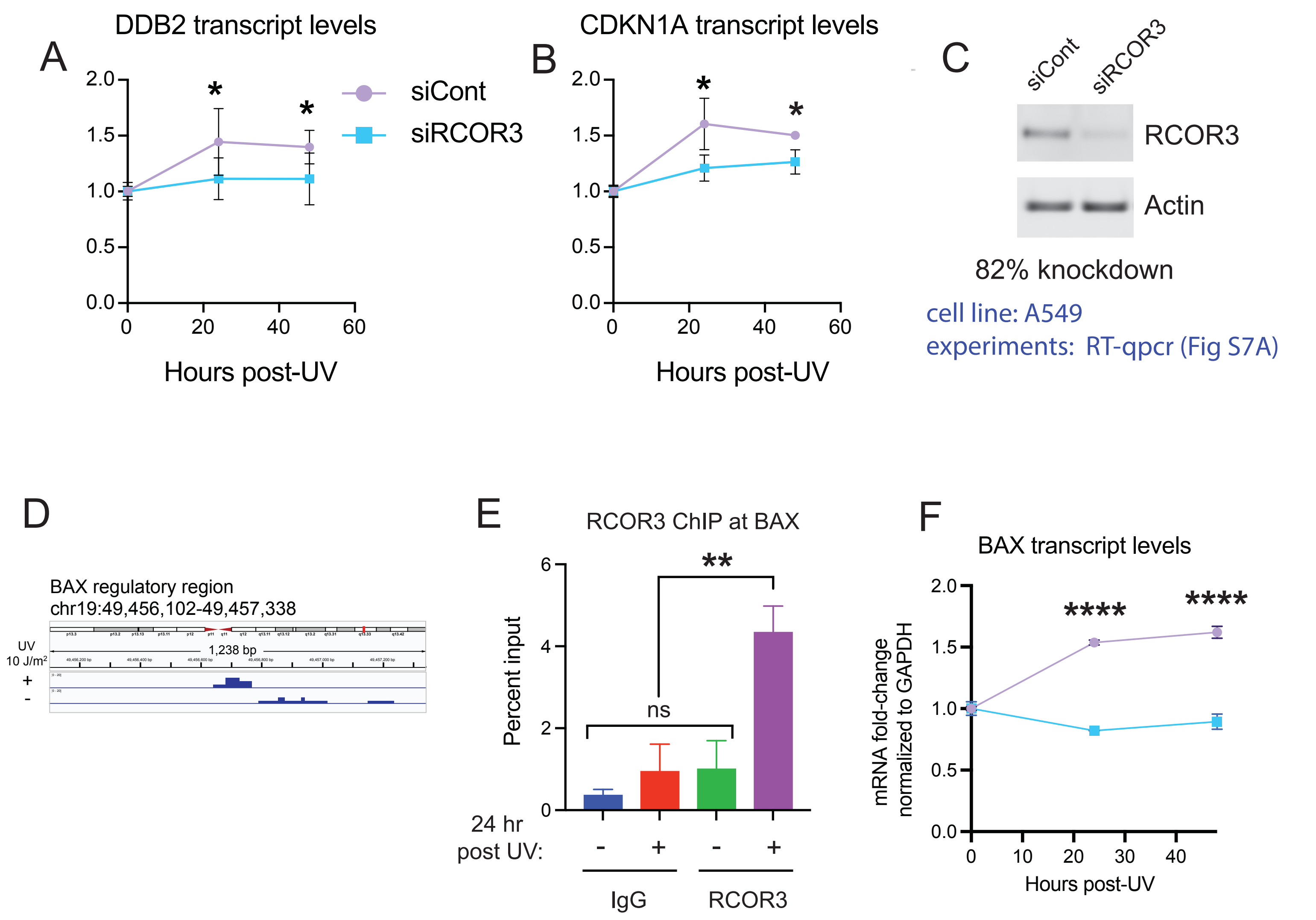

A
DDB2 transcript levels
2.0
1.5
1.0
0.5
0.0
0
20
40
60
Hours post-UV
*
*
siCont
siRCOR3
B
CDKN1A transcript levels
2.0
1.5
1.0
0.5
0.0
0
20
40
60
Hours post-UV
*
*
C
siCont
siRCOR3
RCOR3
Actin
82% knockdown
cell line: A549
experiments: RT-qpcr (Fig S7A)
D
BAX regulatory region
chr19:49,456,102-49,457,338
UV
10 J/m2
+
-
1,238 bp
E
RCOR3 ChIP at BAX
Percent input
6
4
2
0
**
ns
24 hr
post UV:
-
+
-
+
IgG
RCOR3
F
BAX transcript levels
mRNA fold-change
normalized to GAPDH
2.0
1.5
1.0
0.5
0.0
0
10
20
30
40
Hours post-UV
****
****

**Supplementary Figure 6. RCOR dependence of UV-induction of transcripts.** A-B. Quantitation of DDB2 (A) and CDKN1A (B) transcript levels following 10 J/$m^2$ UV at the indicated times in A549 cells with optional depletion of RCOR3 by siRNA. C. Western blot of RCOR3 in A549 cells expressing CRISPRi with either NTC or RCOR3. Percent knockdown is calculated by normalization to beta-actin. D. Genome browser view of G4 peaks at 0 (-) and 24 hr (+) following 10 J/$m^2$ UV at the BAX regulatory region on chr 19. E. RCOR3 occupancy by ChIP at the BAX regulatory region on chr19. F. Quantitation of BAX transcript levels at the indicated timepoints in A375 cells with optional depletion of RCOR3. (IGV) NTC, non-targeting control. IGV, Integrative Genomics Viewer. Chr, chromosome. *, $p < 0.05$; **, $p < 0.01$; ***, $p < 0.001$;****, $p < 0.0001$ (One-way ANOVA for multiple comparisons).

### Figure S7

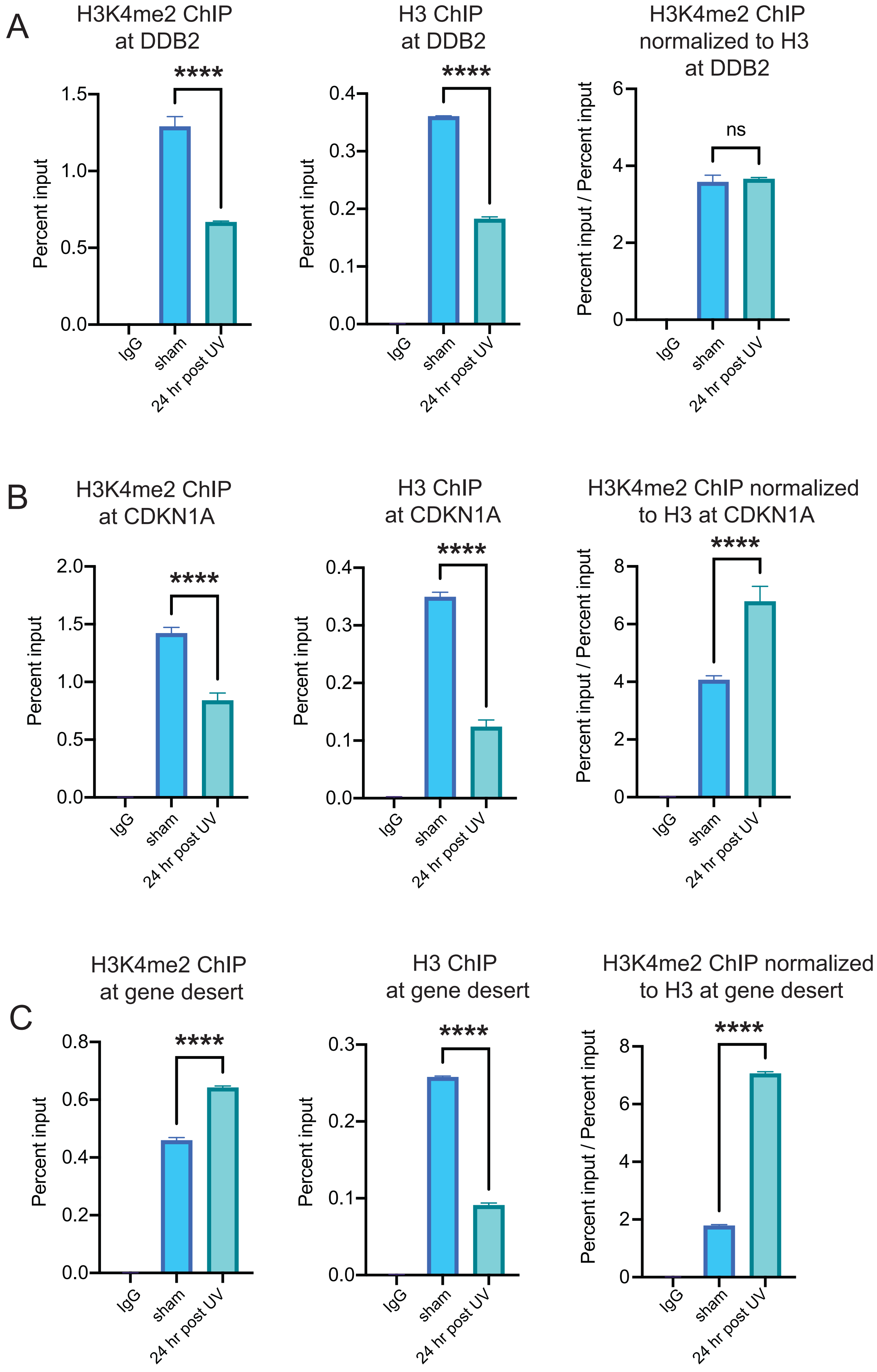
A
H3K4me2 ChIP at DDB2
H3 ChIP at DDB2
H3K4me2 ChIP normalized to H3 at DDB2
B
H3K4me2 ChIP at CDKN1A
H3 ChIP at CDKN1A
H3K4me2 ChIP normalized to H3 at CDKN1A
C
H3K4me2 ChIP at gene desert
H3 ChIP at gene desert
H3K4me2 ChIP normalized to H3 at gene desert
Percent input
Percent input / Percent input
IgG
sham
24 hr post UV
****
ns

**Supplementary Figure 7.** A-C. H3K4me2 occupancy by ChIP at DDB2 (A), CDKN1A (B), and chr4 gene desert (C). chr, chromosome. *, $p < 0.05$; **, $p < 0.01$; ***, $p < 0.001$;****, $p < 0.0001$ (One-way ANOVA for multiple comparisons).

**Supplementary Table Legends**

Table S1. Proteomics data from mTurbo and mTurbo-G4P capture.

Table S2. siRNA oligonucleotide sequences.

Table S3 - Primer sequences for cloning.

Table S4 - gRNA sequences for CRISPRi.

Table S5 - Universal i5 and i7 primers for CUT&Tag.

Table S6 - RT-qPCR primer sequences.

Table S7 - ChIP-qPCR primer sequences.

Table S8 - ProtFiler positive and negative terms.
